# Protracted Functional and Structural Reorganization of Human Prefrontal Cortex Supports Lateralized Category Geometries

**DOI:** 10.64898/2026.05.04.721346

**Authors:** Priscilla Louis, Jewelia K. Yao, Zachariya Yazdani, Ruggaya Musa, Timothy J. Buschman, Jesse Gomez

## Abstract

Despite being visually responsive, how prefrontal cortex represents visual categories across development and how this compares to posterior visual cortex remains unknown. Here, we show that ventrolateral prefrontal cortex (VLPFC) follows a fundamentally distinct developmental trajectory. Using functional and quantitative MRI in children and adults, we demonstrate that VLPFC exhibits reliable and structured category-selective responses in childhood that undergo prolonged reorganization into mature, hemisphere-specific representational geometries. Right VLPFC progressively organizes categories along an animate–inanimate axis, while left VLPFC consolidates a word-selective organization, a lateralization that strengthens across development. This restructuring contrasts sharply with ventral temporal cortex (VTC), which exhibits bilaterally organized, adult-like category structure in childhood. Critically, prefrontal representational structure emerges in alignment with conserved folding patterns of the precentral and inferior frontal sulci, suggesting that cortical anatomy provides a stable scaffold for the maturation of abstract representations. Quantitative MRI further reveals a non-linear trajectory of tissue development in right VLPFC, marked by adolescent reductions in macromolecular content and increased microstructural heterogeneity in adulthood, a pattern not observed in VTC. Together, these findings demonstrate that VLPFC undergoes prolonged, anatomically constrained refinement of category-related geometry characterized by hemisphere-specific taxonomies, rather than simply inheriting representational structure from sensory cortex.

- Unlike VTC (which shows early, bilateral, and stable category organization), VLPFC contains reliable, category-selective representations from childhood that undergo prolonged, hemisphere-specific developmental refinement rather than simple amplification
- Right VLPFC progressively consolidates an animate–inanimate representational axis across development, while left VLPFC establishes an early, stable word-selective organization consistent with language network maturation
- Functional boundaries in VLPFC align with conserved intrasulcal folding motifs (*plis de passage*), demonstrating that anatomical scaffolding of category-selective responses extends beyond sensory cortex to prefrontal association cortex
- Right VLPFC undergoes adolescent pruning of cortical tissue microstructure diverging sharply from the continuous tissue proliferation observed in ventral temporal cortex

## Introduction

The human ability to recognize ecologically meaningful stimuli such as faces^1–3^ and words^4–6^ depends on specialized neural representations in ventral temporal cortex (VTC), which are shaped by visual experience during childhood^2,3,7–9^. These representations are spatially organized relative to cortical folding^10–12^, follow largely linear developmental trajectories^2,6,13–15^ coupled with growth in underlying cortical tissue structures^16^, and preserve their representational structure while increasing in response magnitude^9,13,17–19^. As a result, VTC has served as a canonical model for how experience shapes both functional and structural organization in human cortex, raising the question of how broadly such principles extend beyond sensory cortex.

Whether such organizational principles extend to higher-order visual representations in prefrontal cortex remains unclear. Unlike sensory cortex, prefrontal cortex supports abstract, flexible representations that guide behavior beyond immediate perceptual input^20^. Classical theories therefore emphasize its role in cognitive control and task execution rather than representational content^21–23^. Yet recent work has identified visually responsive regions for faces^27–29^ and words^4,30–32^ within ventrolateral prefrontal cortex (VLPFC), raising the possibility that prefrontal cortex may itself house structured visual representations organized along lateralized axes, analogous to hemispheric specialization for language^24–26^, albeit at a more abstract level than those found in VTC.

How such representations emerge across development is unresolved. Do prefrontal visual responses simply recapitulate organizational principles established in sensory cortex, or do they undergo a distinct and protracted maturation reflecting the extended development of higher-order cognition? Here, we combine functional MRI, quantitative MRI, and developmental gene expression analyses to examine how visual category representations in VLPFC emerge and reorganize across development, using VTC as an early-maturing benchmark. Studying children (ages 5–12) and adults, we find that while VTC exhibits adult-like representational structure early in life, VLPFC follows a prolonged asymmetric trajectory: childhood category responses reorganize into lateralized axes in adulthood, with right VLPFC structured by an animate–inanimate distinction and left VLPFC by a word-selective axis. This functional reorganization aligns with conserved sulcal folding motifs and is accompanied by nonlinear cortical tissue changes, revealing that higher-order visual representations in human PFC emerge through a developmental process coupling late-stage functional reorganization with fine-scale structural remodeling.

## Results

### Category-Evoked Activations in PFC Emerge Early and Sustain into Adulthood

The goal of the present analyses was to characterize how category representations in ventrolateral prefrontal cortex (VLPFC) are organized across hemispheres and how this organization emerges over development. We asked whether VLPFC exhibits systematic category structure, whether this structure is robust within individuals and generalizable across brains, and how its developmental trajectory compares to ventral temporal cortex (VTC), where category organization is established early and bilaterally. Thirty children (ages 5–12, mean 9.17 ± 2.20, 18 females) and thirty young adults (ages 20–29, mean 23.80 ± 1.79, 22 females) completed a functional MRI visual category localizer containing faces, pseudowords, bodies, objects, and places across three runs while fixating and performing an odd-ball detection task^48^ (see **Methods**). A broad region of interest (ROI) encompassing the precentral sulcus (PCS) and inferior frontal sulcus (IFS), where category-selective regions have been previously reported in adults^48^, assessed whether visual responses are present across development, independent of fine-grained functional boundaries.

Both children and adults showed reliable, category-selective responses across all five stimulus categories (all t ≥ 3.50, all FDR ≤ 0.0015), such that a significant proportion of vertices exceeded a selectivity threshold (t-values > 2.5; **Fig. 1A–B)**, indicating that VLPFC represents multiple visual domains from early in development. Overall response magnitude did not differ between children and adults in either hemisphere (LH: p = 0.314; RH: p = 0.303), suggesting that development is not expressed as a global increase in activation. Instead, development was reflected in the redistribution of category-selective territory within VLPFC. In the left hemisphere, word-selective responses expanded by 28.3% and object-selective responses decreased by 39.1% from childhood to adulthood, yielding strong adult dominance of word stimuli by surface area (ANOVA main effect of category: F(4,290) = 7.739, p < 0.001, partial η² = 0.097). In the right hemisphere, face-selective responses increased markedly (59.2%) alongside a comparable reduction in object-selective territory (38.6%), with a more even distribution across categories in adulthood (Effect of category: F(4,290) = 0.562, p = 0.691, partial η² = 0.008). Together, these results demonstrate that VLPFC is robustly responsive to visual stimuli in childhood, with development characterized by the reorganization of category-selective representations rather than broad increases in activation amplitudes.

**Figure 1.**
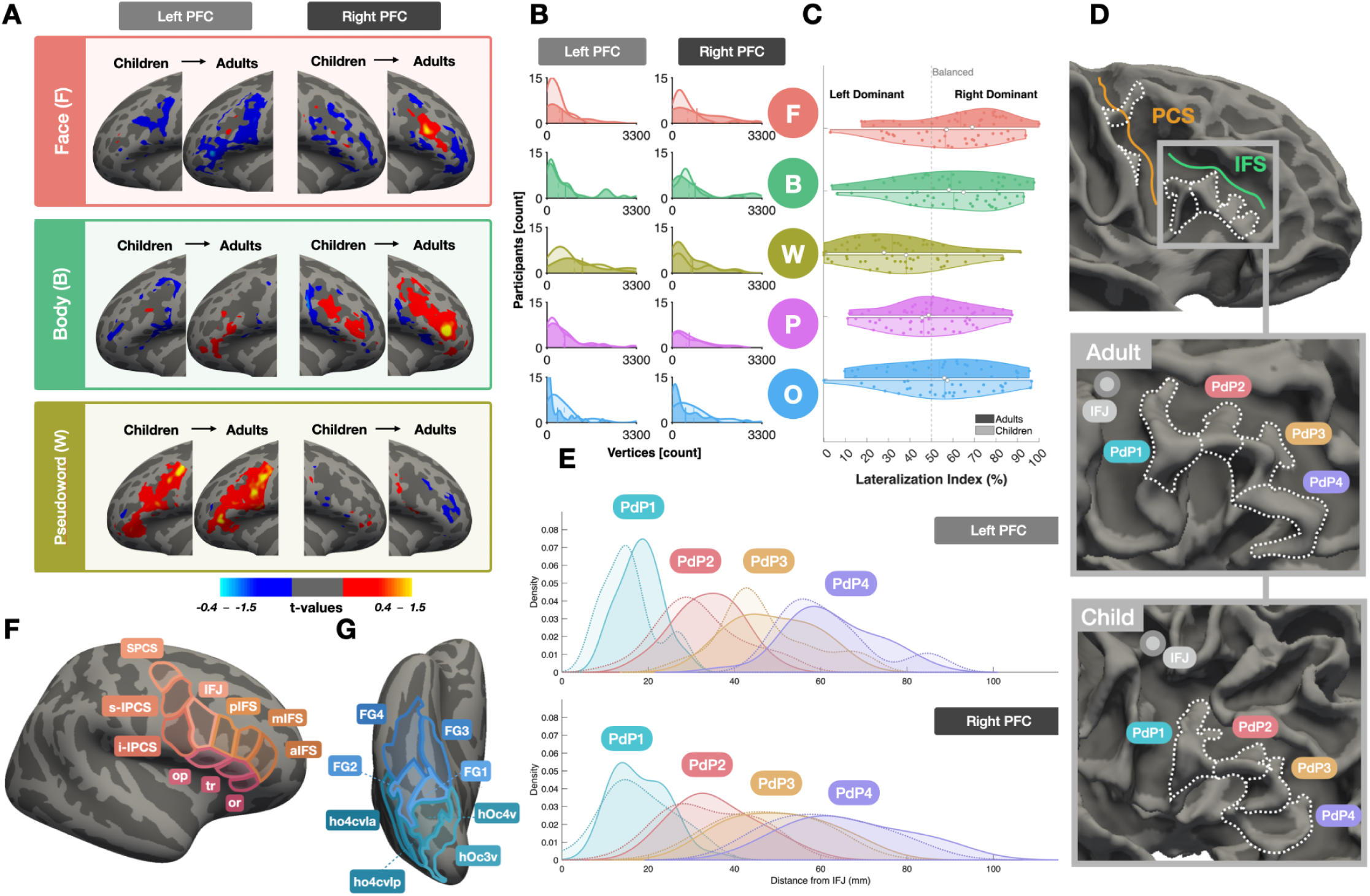
VLPFC shows visual responses and consistent folding patterns across development. **(A)** Average cortical surface maps of category selectivity (preferred > all others) for faces, bodies, and pseudowords on the fsaverage template. Columns show a sample stimulus, left hemisphere maps and right hemisphere maps. Each hemisphere panel shows children (left) and adults (right). Diverging colors indicate positive (red–yellow) and negative (blue–light blue) selectivity. **(B)** Density plots of suprathreshold vertices counts (t-values > 2.5) for five visual categories in each hemisphere. Children are shown with translucent curves and adults with solid curves; vertical lines show group means. **(C)** Split-violin plots of hemispheric dominance (% RH activation relative to LH + RH) per category for children (lighter) and adults (darker). Values >50% indicate RH dominance, <50% indicate LH dominance. **(D)** IFS pleats shown on fsaverage, a sample adult, and a sample child. PdP1 (blue), PdP2 (pink), PdP3 (yellow), PdP4 (purple) are demarcated by white dotted lines; grey circle marks IFJ. **(E)** Geodesic distance distributions from IFj to PdP1–4 for children (dashed) and adults (solid) in left and right hemispheres; colored circles denote PdP anatomical positions. **(F)** Parcellation of VLPFC into 10 ROIs. IFJ: inferior frontal junction, pIFS: posterior inferior frontal sulcus, mIFS: middle inferior frontal sulcus, aIFS: anterior inferior frontal sulcus, SPCS: superior precentral sulcus, s-IPCS: superior component of inferior precentral sulcus, i-IPCS: inferior component of inferior precentral sulcus, op: pars opercularis, tr: pars triangular, or: pars orbitalis. **(G)** Parcellation of VTC into 8 ROIs from Rosenke et al. 2018^62^. Example, open-source stimuli are derived from the fLoc functional localizer package by Stigliani et al 2015 https://github.com/VPNL/fLoc

### Category-Specific Lateralization in Prefrontal Cortex is Developmentally Invariant

While hemispheric lateralization for faces and words has been previously established in VTC^13,16,49–53^, it is unclear whether similar asymmetries extend to prefrontal cortex beyond the known left-lateralization of language^54^, or how they develop. To investigate, we computed a laterality index (LI) reflecting the proportion of visually driven activity (t-values > 2.5) in the right hemisphere relative to both hemispheres **(Fig. 1C)**.

Pseudowords were consistently left-lateralized, whereas faces, bodies, and objects were right-lateralized, with pseudowords showing the strongest leftward bias (adults Cohen’s *d* = 0.90–1.40, children *d* = 0.78–0.98). Critically, this pattern was stable across age groups (all p > 0.05, |*d*| ≤ 0.31), with pseudowords distinct from faces and bodies in both groups (all FDR < 0.05), indicating that category-specific lateralization emerges early and persists throughout development. Thus, VLPFC exhibits an early-emerging and developmentally invariant lateralized organization that parallels hemispheric specialization in ventral visual scream.

### VLPFC Sulcal Landmarks are Anatomically Stable Across Development

Having established VLPFC as visually responsive, we next asked whether category-selective response topographies in VLPFC are spatially stable across development. In VTC, category-selective regions are reliably localized relative to cortical folds, but whether this phenomenon extends to higher-level association cortex in the frontal lobe is unknown.

Many functional responses in VLPFC are buried within precentral and inferior frontal sulci, where intrasulcal folds, known as *“Plis de passage” (PdP)*, create consistent anatomical landmarks. First described over 150 years ago^55^, these sulcal patterns are known to be consistent in adults^57,58^ but their spatial stability across development has not been established. To address this gap, we mapped *PdP* anatomy within the PCS and IFS, building on prior work in occipitotemporal cortex linking functional borders to cortical folds^11,55,56,59,60^, and identified six total *PdPs* reliably observed in both children and adults **(Fig. 1D)**. Two *PdPs* punctuate the PCS: a superior pleat just beneath the precentral-superior frontal junction demarcating the superior and inferior PCS (SPCS, IPCS)^61^, and an inferior pleat just beneath the inferior frontal junction (IFJ) that further bisects the IPCS into its superior and inferior components (s-IPCS, i-IPCS)^59^. Four additional *PdPs* line the inferior bank of the IFS in a spatially consistent, zipper-like arrangement, a configuration not previously characterized.

Geodesic distances from the crown of each *PdP* to the IFJ confirmed no age-related differences in gyrus position in either hemisphere (LH: F(1,58) = 0.14, p = 0.71; RH: F(1,58) = 0.005, p = 0.94), no age-by-gyrus interactions (all p > 0.23), and no hemispheric differences (three-way repeated-measures: all p > 0.24) **(Fig 1E)**. Rather, repeated measures ANOVA including age, gyrus, and hemisphere revealed a robust main effect of gyrus in both hemispheres (all p < 10^-63^), indicating a highly reproducible spatial ordering of the *PdP*.

Leveraging this anatomical consistency, we parcellated VLPFC into ten compartments **(Fig. 1F)**: three PCS subdivisions (SPCS, s-IPCS, i-IPCS), four IFS subdivisions anchored by each PdP (IFJ, pIFS, mIFS, aIFS), and three inferior frontal gyrus components (operculum, triangularis, orbitalis). This parcellation provides a spatially grounded framework for quantifying developmental changes in prefrontal topography.

### Rightward-biased Restructuring of Prefrontal Category Representations Across Development

While category-selective responses in occipitotemporal visual cortex are largely established in early childhood and remain spatially stable across development^2,13,16^, it is unclear whether prefrontal representations follow a similar trajectory or undergo reorganization. To address this, we benchmarked VLPFC responses against ventral temporal cortex (VTC) using an existing FreeSurfer cytoarchitectonic parcellation of VTC encompassing eight regions across the occipitotemporal sulcus (OTS), fusiform gyrus (FG), collateral sulcus (CoS), and the lingual gyrus^62^. The borders between these regions follow cortical folding^11^ and encompass the relevant extent of high-level visual cortex^15,37^.

We confirmed that category-selective responses were highly consistent across age groups in VTC **(Fig. 2A–B)**, with faces peaking in FG2 and places in FG3 as expected. Correlating mean per-ROI category response vectors between children and adults confirmed near-identical topographies in both hemispheres (RH: r = 0.94, FDR<0.001; LH: r = 0.97, FDR<0.001), establishing VTC as a stable developmental benchmark.

**Figure 2.**
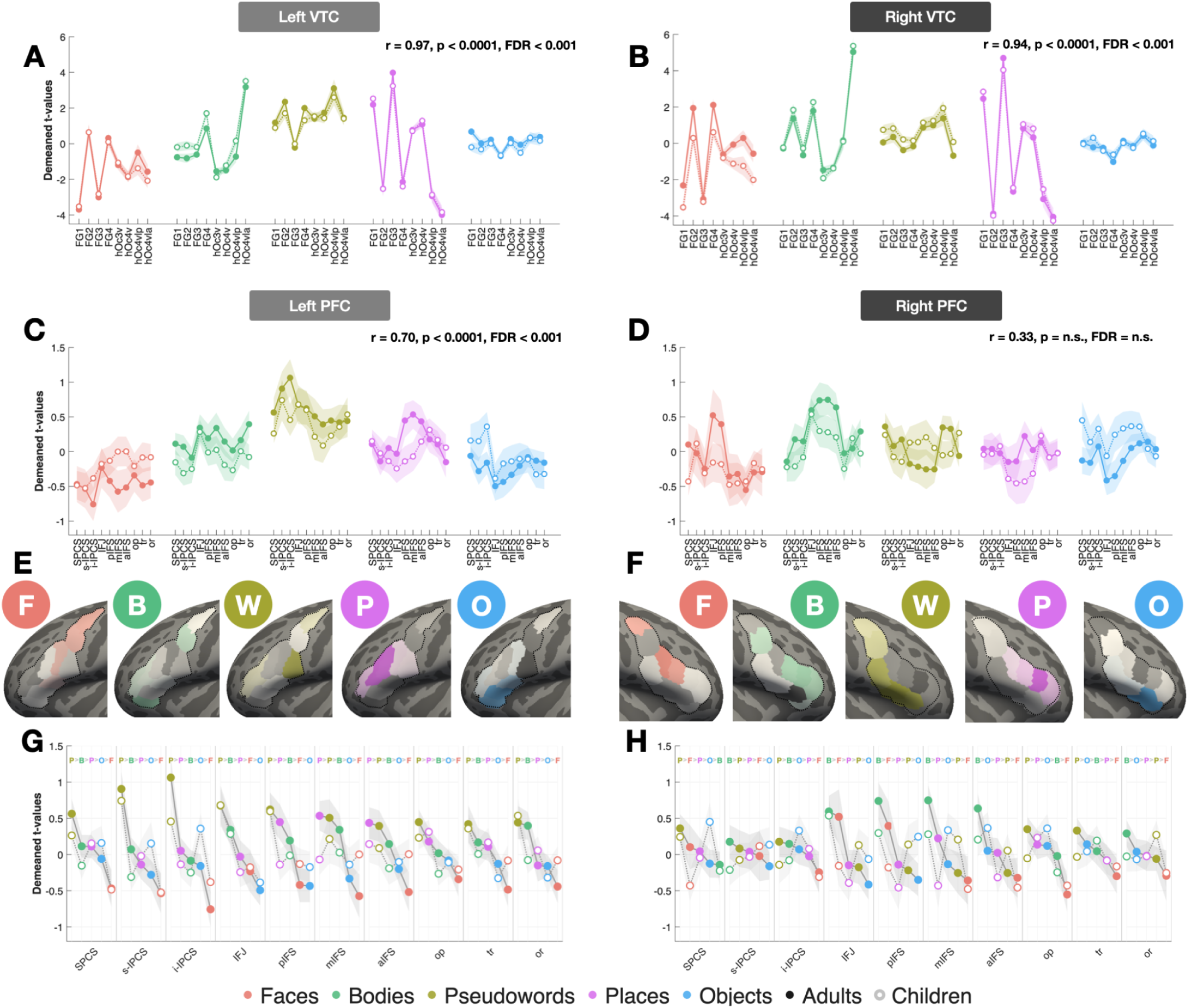
Category response topographies develop more in VLPFC than VTC, particularly in the right hemisphere. **(A)** LH VTC lineplots of z-scored category-selective responses (t-values) grouped by contrast and centered relative to the adult baseline. Solid/dashed lines indicate adults and dashed/open markers indicate children; shading indicates ± SEM. **(B–D)** Same as A, but for RH VTC, LH VLPFC, and RH VLPFC, respectively. **(E)** Z-scored developmental difference maps (adults minus children, normalized within each system) projected onto the LH VLPFC fsaverage surface. A single z-scored value was assigned to each ROI and projected onto cortical surface vertices, resulting in uniform coloring within regions. Positive values indicate stronger responses in adults than children, whereas negative values indicate stronger responses in children than adults. **(F)** Same as E, but for RH PFC. **(G)** LH VLPFC responses grouped by ROI, showing category trajectories ordered from strongest to weakest responses in adults (F=faces, B=bodies, yellow P=pseudowords, purple P=places, O=objects). **(H)** Same as G, but for RH PFC. Example, open-source stimuli are derived from the fLoc functional localizer package by Stigliani et al 2015 https://github.com/VPNL/fLoc

VLPFC showed a starkly different pattern **(Fig. 2C–H)**. Left hemisphere response profiles were significantly correlated across age groups but weaker than VTC (r=0.70, FDR<0.001). However, right hemisphere profiles were not significantly correlated (r = 0.33, FDR=n.s.), a coefficient significantly smaller than the corresponding VTC correlation (z-score = 6.57, p < 0.001). This was reflected at the level of individual ROIs: for every visual category in the right hemisphere, the region showing peak selectivity in adults differed from that in children, and no ROI preserved its adult stimulus preference ranking in children **(Fig. 2G–H)**. A comparable pattern was observed in the left hemisphere, including pseudowords which peaked in i-PCS in adults but s-IPCS in children. Consistent with this hemispheric asymmetry, interhemispheric response profile correlations were lower in VLPFC than VTC for both children (Fisher z-transform = 0.594 vs 1.208) and adults (Fisher z-transform = 0.668 vs 1.062), highlighting that VLPFC is less bilaterally coordinated. Together, these results demonstrate that VLPFC category representations undergo pronounced developmental reorganization, particularly in the right hemisphere, rather than the stable spatial scaling observed in VTC.

### Cross-Validated Reliability Reveals Hemisphere-Specific Taxonomies in VLPFC

A central question following the observed developmental reorganization of VLPFC is whether children’s VLPFC responses reflect reliable, idiosyncratic representations or simply noise. VTC responses in the same participants are largely stable from childhood to adulthood, demonstrating that adult-like patterns can indeed be measured in children. We therefore ask if the developmental topography in PFC is similarly reliable, and if so, whether they reflect fundamental changes in the way information is represented across development.

Using leave-one-run-out (LORO) cross-validation **(Fig. 3A,E)**, we computed voxelwise pattern similarity (Pearson correlation) across all voxels in VLPFC. Within-category correlations were calculated by correlating the pattern for the same category across independent runs, while between-category correlations were calculated by correlating patterns of different categories across runs. Across hemispheres and age groups, within-category correlations consistently exceeded between-category correlations, confirming reliable VLPFC selectivity profiles. In right VLPFC, within-category correlations significantly exceeded between-category correlations in both children (r = 0.089 ± 0.017 vs –0.022 ± 0.004; t(29) = 5.182, p = 1.53e-05, *dz* = 0.946) and adults (r = 0.071 ± 0.017 vs –0.017 ± 0.004; t(29) = 4.103, p = 0.000302, *dz* = 0.749), with comparable reliability in left VLPFC. VTC exhibited robust within-category reliability bilaterally **(Fig. S3)**, consistent with prior work. These results confirm that VLPFC representations are stable and functionally organized.

**Figure 3.**
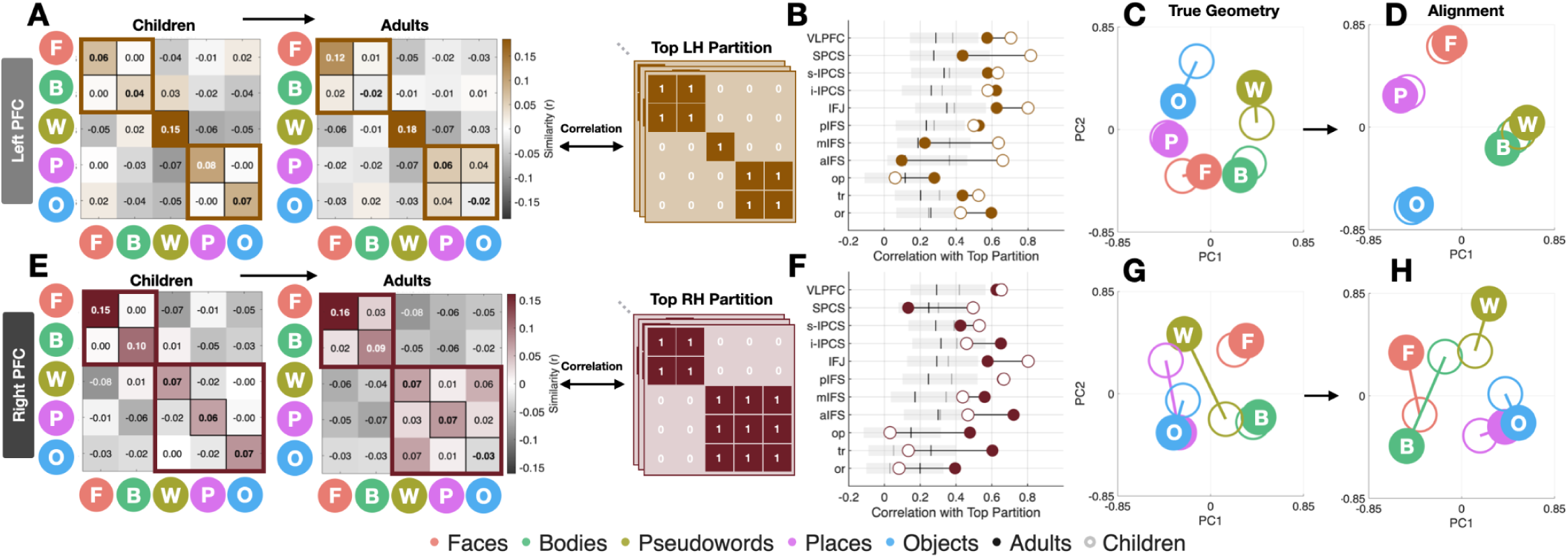
Representational geometry and clustering of category-selective responses in VLPFC across development. **(A)** LH VLPFC representational similarity matrices (mean pairwise Pearson correlation between voxel-wise t-value maps for five categories) computed using leave-one-run-out cross-validation, shown separately for children and adults. Diagonal values index within-category reliability; off-diagonal values index cross-category similarity. Colorscale is centered at zero. **(B)** LH VLPFC dumbbell plots comparing neural correlations with all binary partition models for adults (filled) and children (open). Highlighted partitions reflect the best-fitting organizational schemes: FB_W_PO (animate / pseudoword / spatial-inanimate) for LH and FB_WPO (animate / inanimate for RH). Gray shading indicates variability across all remaining partitions, and vertical lines show group-specific mean correlations. **(C)** PCA of vertex-wise category-selective response patterns in LH, plotted in PC1–PC2 space (z-scored across vertices). Adult maps are filled and child maps are open. Lines connect matched categories across age groups. **(D)** Same as C, following Procrustes alignment of children to the adult configuration. **(E–H)** Same as A–D, but for RH PFC. Example, open-source stimuli are derived from the fLoc functional localizer package by Stigliani et al 2015 https://github.com/VPNL/fLoc

We next tested whether VLPFC similarity matrices exhibit coherent higher-order structure, guided by the animate–inanimate organization that robustly characterizes VTC^63–65^. In right VLPFC, adults showed a clear animate–inanimate block structure (mean within-block r = 0.047 ± 0.010 vs. between-block r = –0.052 ± 0.011; t(29) = 4.544, p = 8.98e-05, *dz* = 0.830), which was absent in left VLPFC **(Fig. 3A,E).** Children already showed early animate–inanimate segregation in right VLPFC (within-block r= 0.030 ± 0.009 vs. between-block r = –0.033 ± 0.009; t(29) = 3.548, p = 0.00134, *dz* = 0.648), indicating protracted rather than abrupt development. Left VLPFC instead showed a tripartite semantic blocking which divided animate (face, body), language (pseudoword), and spatial (place, object) categories in both adults (mean within-block r = 0.050 ± 0.012 vs. mean between-block r = –0.026 ± 0.007; t(29) = 4.060, p = 0.00034, *dz* = 0.741) and children (mean within-block r = 0.043 ± 0.010, mean between-block r = -0.023 ± 0.005; t(29) = 4.451, p = 0.000116, *dz* = 0.813), suggesting that this left hemisphere organization is established early (**Fig 3B,F**).

To confirm that these structures reflect principled representational taxonomy rather than an arbitrary grouping, we conducted an algorithmic splitting approach testing all 75 unique ways of dividing the five categories into two non-overlapping groups (excluding duplicates). Each partition was encoded as a binary model and correlated with cross-validated LORO similarity matrices **(Fig. 3B,F)**. In right VLPFC, the animate–inanimate partition ranked first in adults (r = 0.815; rank = 1/75; percentile = 0.013) and within the top quartile in children (r = 0.593; rank = 18/75; percentile = 0.240), confirming that this structure is present early and sharpens with development. In left VLPFC, a word-selective partition ranked first in adults (r = 0.658; rank = 1/75; percentile = 0.013) and remained highly ranked in children (r = 0.644; rank = 11/75; percentile = 0.147), indicating that left-hemisphere organization is established early but continues to refine across development.

### Bilateral Semantic Organization in VLPFC Emerges Through Right-Hemisphere Maturation

Having established reliable category selectivity, we next examined how category responses are geometrically organized across development using principal component analysis (PCA), where distances between categories reflect similarity of neural responses. We compared children and adults before (**Fig. 3C–D)** and after Procrustes alignment (**Fig. 3G–H**). Pre-alignment geometries characterize the internal representational structure within each group, allowing us to assess whether children exhibit coherent structure independent of their relationship to adults. Post-alignment quantified how well children’s category structure aligns with adults while preserving the relative distances among categories.

Prior to alignment, category responses formed a structured representational space in both groups and hemispheres **(Fig. 3C–D)**. To quantify this structure, we computed pairwise Euclidean distances between category positions in PCA space for each subject. In both hemispheres and age groups, all category pairs were robustly separated from zero (all t > 11.6, all p < 10⁻¹²).

Within-category distances represented the average Euclidean distance among items inside the same category cluster (e.g. distances among Faces only, Bodies only, etc.) and between-category represented the distance between centroids of the semantic clusters (e.g. Animate [Face/Body] centroid distances vs Inanimate [Word/Place/Object] centroids). Adults exhibited robust animate–inanimate separation in both hemispheres (Face/Body vs. Place/Object: RH: (t(29) = 4.17, p = 2.7×10^-77^; LH: t(29) = 3.68, p = 2.87×10^-71^) with children showing a similar but reduced structure (RH: t(29) = 3.75, p = 3.89×10^-72^; LH: t(29) = 3.05, p = 2.32×10⁻⁶²). Critically, pseudoword representations were misaligned in children: word–body distances were significantly smaller than animate–inanimate axis distanced in both hemispheres (RH: t(29) = −4.01, p = 0.00039); LH: (t(29) = −2.69, p = 0.0117), indicating that words are not yet integrated into the dominant representational geometry observed in adults.

After Procrustes alignment, children exhibited a pronounced hemispheric asymmetry in VLPFC **(Fig. 3D,H)**: correlation with a mean-child template, was significantly higher in the left hemisphere than in the right (Paired t-test: LH = 0.697 ± 0.019 vs. RH = 0.561 ± 0.022; *dz* = 0.793, p = 1.56 × 10⁻⁴) with corresponding lower geometric dispersion, defined here as the mean Procrustes distance between each child’s PC coordinates and the child group average (Paired t-test: LH = 0.537 ± 0.030 vs. RH = 0.700 ± 0.026; *dz* = −0.71, p = 5.30 × 10⁻⁴), Adults showed no hemispheric differences in either metric (similarity: *dz* = 0.046, p = 0.081; dispersion: p = 0.67), indicating that mature VLPFC organization is bilaterally balanced. Cross-age alignment showed no hemispheric asymmetry (children-to-adult mean: LH = 0.524 ± 0.028 vs. RH = 0.562 ± 0.027; *dz* = 0.083, p = 0.749), suggesting that asymmetry observed in children reflects internal developmental structuring rather than differential proximity to adult templates.

VTC showed a contrasting profile **(Fig S3)**: Children exhibited strong left-hemisphere coherence that persisted into adulthood (children similarity: LH = 0.751 ± 0.025 vs. RH = 0.538 ± 0.026; dz = 1.74, p < 10⁻⁶; adults similarity: LH = 0.731 ± 0.030 vs. RH = 0.759 ± 0.027), indicating stable rather than converging lateralization. Together, VLPFC and VTC show fundamentally distinct developmental trajectories: VLPFC progresses from childhood hemispheric asymmetry toward bilateral balance, while VTC maintains persistent lateralization across development.

### Cross-Subject Decoding Supports Hemisphere-Specific Functional Reorganization

Building on our PCA analyses, we next asked whether VLPFC representational geometry predictively generalizes across individuals, distinguishing shared coding schemes from participant-specific factors (e.g., attentional state or strategy). We implemented a leave-one-subject-out (LOSO) decoding approach **(Fig. 4A–B)**, first testing whether fine-grained category information is decodable across individuals, then whether cluster solutions derived from a group predict the representational organization of a held-out participant. True decoding accuracy was compared against a shuffled baseline, generated by randomly permuting category labels 1000 times, such that true-minus-shuffled accuracy reflects information above chance. Category identity was reliably decodable above chance across both hemispheres and age groups. Left VLPFC showed robust cross-subject decoding (children: 0.353; adults: 0.340; shuffled ∼0.20), while right VLPFC showed weaker but reliable performance (children: 0.273; adults: 0.273; shuffled ∼0.20), consistent with its more protracted representational development.

**Figure 4.**
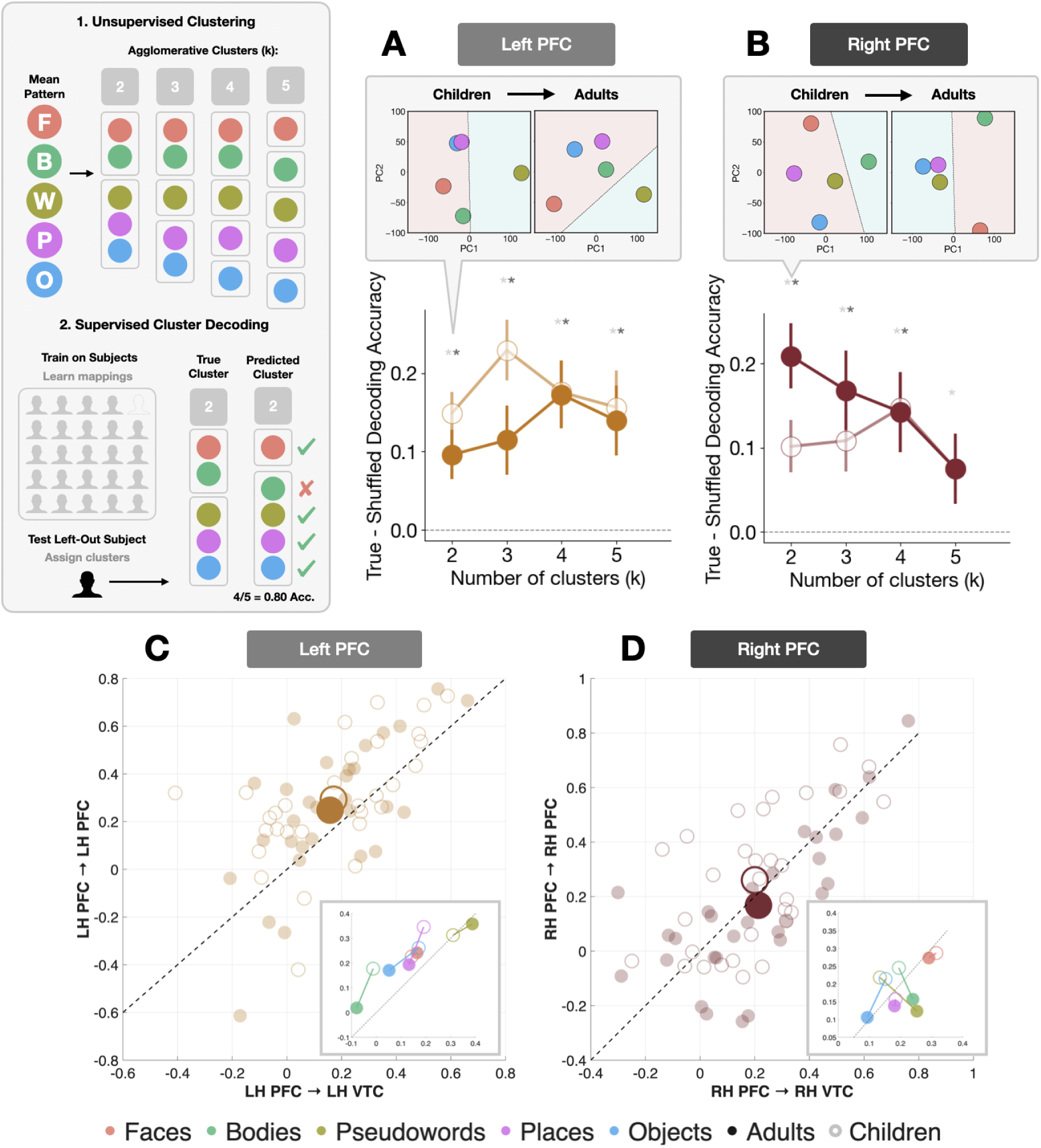
Decoding and representational coupling of category-selective responses in VLPFC across development. **(A)** LH VLPFC true-minus-shuffled LinearSVC decoding accuracy for cluster membership (k = 2–5). Solid lines show the mean across subjects; error bars indicate ± SEM. Adults shown at full opacity, children with reduced opacity. Inset: PCA decision boundary maps for k = 2 clusters in children (left) and adults (right). **(B)** Same as A, but for RH VLPFC. **(C)** LH VLPFC stability–coupling scatterplot: intrinsic VLPFC stability (y-axis) versus PFC–VTC similarity (x-axis). Each point represents an individual participant (children: unfilled circles; adults: filled circles). Large markers denote the group means. The identity line (y = x) indicates equilibrium between internal stability and cross-regional coupling; deviations reflect dominance in either direction. Inset: Category-level mean coordinates for children and adults within the same space. **(D)** Same as C, but for RH VLPFC. Example, open-source stimuli are derived from the fLoc functional localizer package by Stigliani et al 2015 https://github.com/VPNL/fLoc

Higher-order clustering revealed a striking hemispheric asymmetry. In left VLPFC, two-cluster solutions consistently segregated words from all other categories in both children and adults **(Fig. 4A)**, confirming early language-selective organization^66–68^. This partition was robust in children (accuracy = 0.693 ± 0.023 vs. shuffled: 0.544 ± 0.002; true–shuffled: t(29) = 5.375, p < 0.0001, *d* = 0.981) and adults (true–shuffled = 0.0960 ± 0.0321; t(29) = 2.988, p < 0.006, d = 0.545), with the modest adult reduction reflecting increased internal differentiation among non-word categories rather than loss of shared structure. This is consistent with the preserved geometry observed in the Prorustes analysis **(Fig. 3D)**. All cluster solution accuracies (k = 2–5) were significantly above chance in both groups (all p < 0.01; all d > 0.467).

Right VLPFC showed clear developmental change **(Fig. 4B)**. In children, two-cluster solutions separated bodies from all other categories (true–shuffle: 0.1014 ± 0.0316; t(29) = 3.211, p = 0.003), reflecting nascent but incomplete animate–inanimate organization. By adulthood, clusters sharpened into a canonical animate–inanimate structure (accuracy: 0.713; true–shuffle = 0.209 ± 0.0392, p < 0.0001, *d* = 0.972). Two-way ANOVAs revealed no age-by-granularity interactions, indicating that developmental change reflects the emergence of dominant cluster structure rather than sensitivity to the number of clusters tested.

Low-dimensional PCA projections corroborated these patterns, using convex hull analyses to compute the minimal polygon enclosing all categories and thus assessing developmental changes in representational dispersion. Left VLPFC showed modest expansion of representational spread across development (convex hull: children 12,037; adults 13,024), consistent with increasing differentiation among non-word categories within a stable organizational framework. Right VLPFC showed a reduction in representational spread (children: 14,971; adults: 13,456), consistent with the progressive clustering of categories along the animate–inanimate axis. Together, these results demonstrate that VLPFC encodes shared, decodable category representations from childhood, with hemispheric asymmetry in developmental trajectory: Left VLPFC establishes stable language-selective organization early, while right VLPFC undergoes protracted refinement toward an animate–inanimate axis.

### Prefrontal Stability and Posterior Integration is Coordinated Early and Refines with Maturation

The preceding analyses revealed two defining features of prefrontal development: (1) bilateral PFC exhibits stable, distinct semantic geometries within and across subjects, and (2) right PFC shows delayed maturation, with an early animate bias for bodies that grows to include faces. This raises a mechanistic question: What is driving this functional restructuring? Does PFC progressively align with VTC representational geometry across development (*coupling*), or does it consolidate a geometry unique to prefrontal cortex (*stability*)?

To distinguish between these possibilities, we constructed a *coupling–versus—stability* coordinate system (**Figure 4C-D)**. The horizontal axis indexes PFC–VTC *coupling*, computed by correlating the similarity matrix derived from one split of a LORO analysis with the similarity matrix from the other region, and averaging this estimate across all data split combinations. The vertical axis indexed intrinsic PFC *stability*, computed analogously by correlating the similarity matrices from splits of the data in a LORO analysis, comparing PFC to itself across splits (i.e., a split-half reliability metric). Each subject yielded a single (X,Y) coordinate per hemisphere. *Stability* and *coupling* were positively correlated in both children (LH: r = 0.572, p = 9.71 × 10⁻⁴; RH: r = 0.597, p = 5.00 × 10⁻⁴) and adults (LH: r = 0.656, p = 8.20 × 10⁻⁵; RH: r = 0.739, p = 3.00 × 10⁻⁶), indicating that internal stability and long-range coupling are coordinated from early in development and become increasingly aligned with maturation.

Signed distance from the identity line revealed that children were biased toward internal PFC coherence over VTC coupling (t(29) = 3.03, p = 0.005), while adults showed no such offset (t(29) = 0.74, p = 0.465), indicating that maturation reflects a progressive balancing of local stability and long-range integration. This pattern was lateralized (F(1,58) = 7.58, p = 0.008), with no age interaction (all ANOVA p > 0.30). Left VLPFC maintained internal stability dominance across both age groups (children: mean = 0.086, t(29) = 2.927, p = 0.007; adults: mean = 0.064, t(29) = 2.200, p = 0.036), while right VLPFC remained near equilibrium, with adults trending toward VTC coupling dominance (children: mean = 0.043, p = 0.135; adults: mean = –0.032, p = 0.210). Thus, left VLPFC word-related structure is internally dominant and developmentally stable, whereas right VLPFC animate–inanimate organization gradually strengthens its coupling with VTC.

Category-level displacement vectors, which capture the movement in *stability–coupling* space from childhood to adulthood, showed consistently significant magnitude (all magnitude tests p < 0.001) and structured directionality. In left VLPFC, bodies, places, and objects predominantly shifted toward lower coupling and stability, while faces and words shifted toward higher stability and stronger VTC coupling. In right VLPFC, most categories drifted toward relatively greater VTC coupling, with words showing the most pronounced rightward-downward displacement.

### Nonlinear Development of Fine-Scale Tissue Microstructure in Right VLPFC Mirrors Functional Reorganization

The restructuring of VLPFC raises a fundamental question: does the underlying cortical tissue follow a distinct developmental trajectory, potentially supporting these functional changes? We analyzed quantitative MRI (qMRI) data in a lifespan sample (N = 82, ages 5–54 years)^68^, focusing on R1 (sensitive to tissue composition and myelin) and macromolecular tissue volume (MTV, reflecting tissue density), examining both VLPFC and VTC to test whether their divergent functional trajectories entail distinct tissue development.

VTC showed steady increases in both R1 **(Fig. 5A)** and MTV **(Fig. 5B)** from childhood through adulthood, indicative of continuous macromolecular proliferation, suggesting water in the cortex is being continuously displaced by macromolecules including myelin. Right VLPFC followed a distinct and nonlinear trajectory: tissue properties increased from childhood into adolescence, dipped during late adolescence, then recovered in adulthood **(Fig. 5A-B)**. This adolescent dip was absent in left VLPFC, where both R1 and MTV increased steadily from adolescence onward. A cortical lobe x age interaction confirmed divergent temporal and prefrontal tissue trajectories (F(1,29) = 6.18, p < 0.001), and a hemisphere x age interaction within VLPFC confirmed divergent left–right maturation (F(1,29) = 5.64, p = 0.02). To replicate this observation, the same children (N = 25) and young adults (N = 20) who completed the aforementioned fMRI experiments completed quantitative imaging. Right VLPFC showed statistically equivalent R1 and MTV across both age groups (R1: t(1,40.23) = −0.66, p = 0.51; MTV: t(1,38.68) = −1.54, p = 0.13), consistent with transient adolescent tissue loss followed by recovery rather than continuous growth. Thus, two separate datasets confirm that right prefrontal cortical tissue shows a unique developmental trajectory.

**Figure 5.**
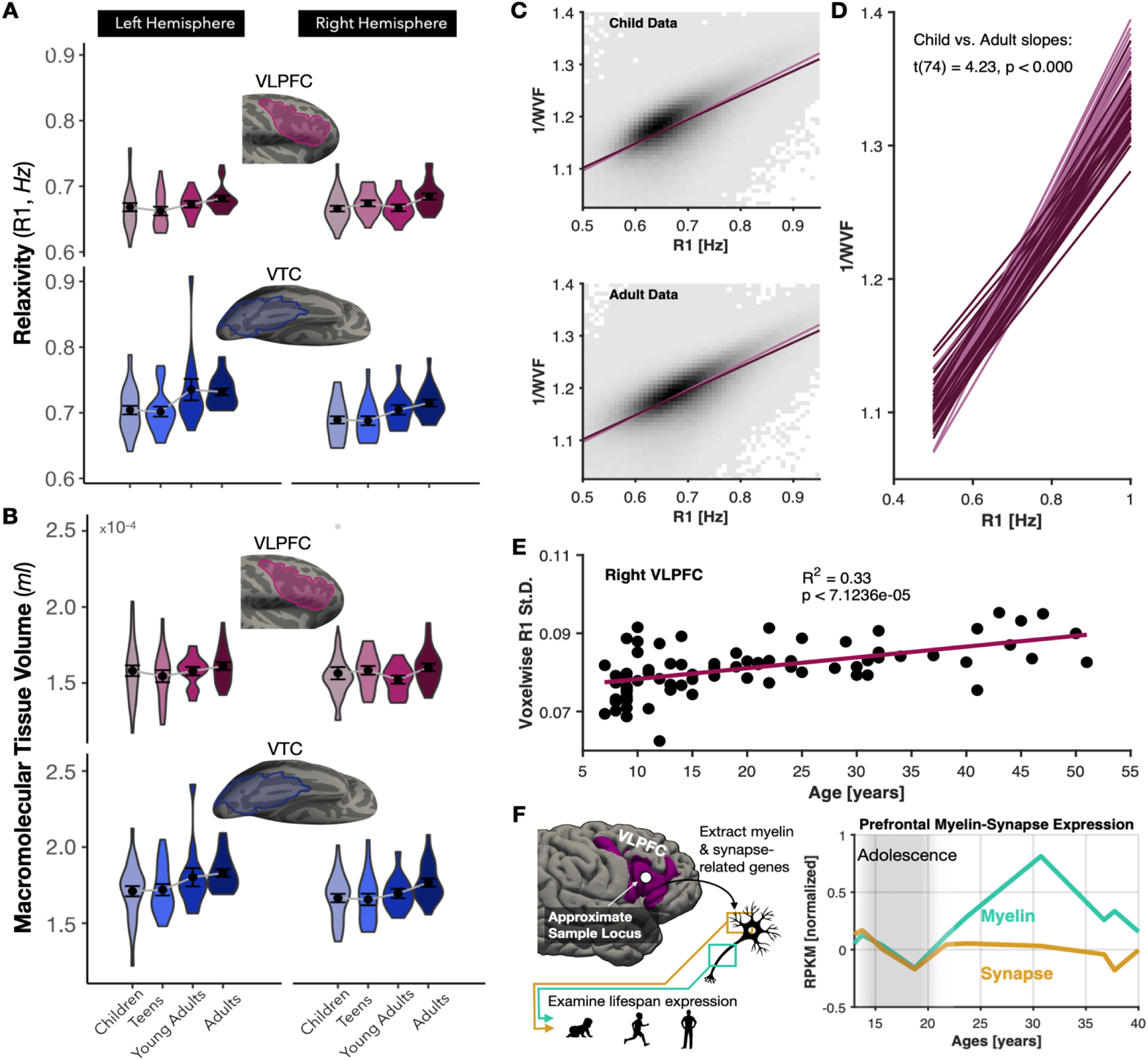
Tissue development diverges between prefrontal and temporal cortex. **(A)** Violin plots of relaxivity (R1) values for children (ages 5 -10), teens (ages 11 - 17), young adults (ages 18 - 24, blue), and adults (ages 25 - 54) in left and right VLPFC (top) and VTC (bottom). Inset: cortical surfaces with TOI outlines. **(B)** Same as A for MTV. **(E)** Scatterplot of R1 standard deviation across VLPFC versus age, with linear fit (cyan; R^2^ and p inset). **(C)** Binned voxelwise scatterplots of R1 versus inverse water volume fraction (1/WVF) pooled across child (top) and adult (bottom) VLPFC, with linear fit for children (light purple) and adults (dark purple) overlaid for comparison. **(D)** Per-subject linear fits of R1 versus 1/WVF in VLPFC. Inset text shows two-sample t-test comparing the slopes between children and adults. **(F)** Developmental gene expression in VLPFC for myelin-associated (cyan) and synapse-associated (gold) gene sets, normalized to mean adolescent expression (gray shaded region, ages 11-21). Donors aged 11-40; expression in RPKM. Inset brain shows tissue sampling location.

To characterize the physicochemical nature of this trajectory, we examined the R1–MTV relationship, where deviations from linearity indicate the presence of hydrophobic materials altering proton relaxation. Children showed a steeper, more constrained R1–MTV slope than adults **(Fig 5C–D)**, and inter-voxel R1 standard deviation increased linearly with age **(Fig 5E)**, indicating progressive variegation of tissue composition across VLPFC. Together, these metrics suggest that in adulthood, a given difference in tissue volume yields a disproportionately larger R1 difference than in childhood, which is consistent with a cortical environment increasingly biased toward myelinated structures.

To test whether adolescent pruning selectively excised non-myelinated tissue, we examined developmental gene expression in VLPFC from a development gene expression dataset^71^, comparing gene families associated with myelin versus synaptic structures^72^. Both gene sets declined during adolescence. However, myelin-associated gene expression recovered into adulthood while synaptic gene expression remained at post-adolescent pruning levels **(Fig 5F)**. This pattern corroborated the qMRI findings suggesting that adolescent pruning disproportionately removes non-myelinated structures like synapses or dendrites, leaving an adult VLPFC tissue environment relatively enriched for myelin.

## Discussion

Traditionally, prefrontal cortex has been conceptualized as a bidirectional hub that integrates context, rules, and task-relevant signals, modulating domain-specific sensory areas while also maintaining abstract, amodal representations^21,74,75^. In this framework, PFC is often viewed as primarily shaping perceptual computations in posterior cortex through attention and top-down control rather than retaining perceptual information itself. While domain-specific patches have been noted, such as language-related clusters in left LPFC or face-selective patches in right IFJ, these have been previously interpreted as exceptions embedded within a largely amodal workspace^72,76–84^.

The present study updates these classical accounts. We show that the category representational architecture of VLPFC undergoes prolonged, hemisphere-specific developmental refinement. While children already exhibit category-selective responses, the spatial topography and multivariate geometry of these responses differ markedly from adults relative to conserved intrasulcal folding motifs. Across development, these representations reorganized into structured, lateralized geometries: right VLPFC progressively consolidates an animate–inanimate axis, while left VLPFC establishes a word-selective organization consistent with its role in language systems. This reorganization is most striking in the right hemisphere, where childhood response topographies are reliable within individuals but uncorrelated with adult spatial patterns. Critically, this functional restructuring is accompanied by adolescent pruning of cortical tissue in right VLPFC, diverging sharply from the continuous tissue proliferation observed in VTC.

### Representational Structure versus Domain-General Control

Our findings are inconsistent with domain-general control signals, low-level stimulus properties, or task-state fluctuations. Stimuli were carefully matched on low-level visual features^73^, participants performed a single stable oddball task across runs, and category-selective patterns were reproducible within and across individuals. If responses reflected task-state variability, similarity structure would not replicate across runs or generalize across subjects. Instead, convergent evidence from activation profiles **(Fig. 2)**, cross-validation similarity matrices **(Fig. 3A–B,E–F)**, PCA geometry **(Fig. 3C–D,G–H)**, and cluster decoding **(Fig. 4A–B)** consistently demonstrates high-order representational organization that is internally reliable and population-reproducible. Benchmarking against VTC further confirms that VLPFC does not duplicate sensory representations but transforms and reorganizes categorical information.

### Hemispheric Specialization

Left VLPFC shows early consolidation of word-selective organization, consistent with the rapid maturation of left perisylvian language networks that increasingly constrain left VLPFC toward linguistic semantics^73,85–95^. Cross-validated decoding confirmed that word-isolating partitions outperform nearly all alternatives from childhood onward **(Fig. 4A)**, suggesting that left VLPFC provides a scaffold for linguistic semantic representation. This early emerging left-hemisphere commitment leaves right VLPFC relatively unconstrained and available for representing higher-level visual–semantic distinctions. Right VLPFC undergoes prolonged restructuring along an animate–inanimate axis, likely shaped by convergent ventral visual and posterior parietal inputs^87,97^, early experiences linking animacy with agency, and the functional demand to supply the left hemisphere with a semantic prior that interfaces with emerging lexical systems. This potentially also lays an early foundation for right VLPFC roles in error monitoring, inhibitory control, and visuospatial reasoning^85,92,94,96,97^.

### Comparison with VTC

VLPFC and VTC follow fundamentally distinct developmental trajectories. VTC exhibits early-emerging and bilaterally organized animate–inanimate structure, whereas VLPFC undergoes extended asymmetric refinement, particularly in the right hemisphere. This distinction indicates that prefrontal category representations do not simply recapitulate the organizational principles of sensory cortex but instead reflects a prolonged developmental regime. Prior work observed recycling of body representations for faces and words in VTC during childhood^13^. By contrast, we observe bilateral increases in body representation along the IFS in prefrontal cortex, further distinguishing these developmental trajectories. Multivariate geometry analyses reveal a corresponding double dissociation: childhood VLPFC is characterized by left-dominant hemispheric asymmetry, while adulthood is associated with improved interhemispheric balance; this is the inverse of VTC, which maintains a stable left-lateralized coherence across development.

### Posterior-Anterior Coupling

Intrinsic prefrontal representational stability was positively correlated with PFC–VTC similarity in both age groups, with this coordination strengthening across development **(Fig. 4C–D)**. Children showed a bias toward intrinsic prefrontal coherence while adults showed greater balance between internal stability and cross-regional similarity. We interpret this not as PFC duplicating VTC content, but as progressive tuning of prefrontal representations to be better coordinated with an already-mature posterior geometry. Supporting this, an adult-like animate–inanimate boundary is present in VTC but absent in right VLPFC children. Thus, maturation involves rebalancing of prefrontal representational manifolds toward increased face representation at the expense of objects. However, this interpretation remains correlational, and causal relationships await future investigation.

### Anatomical Scaffolding

A novel finding is that functional boundaries in VLPFC align with conserved intrasulcal pleating motifs within PCS and IFS, called *plis de passage (PdP)*^55,56,59,60,98^. Prior studies defined a connection between function and structure in PFC in terms of its rostro-caudal and dorso-ventral gradients^86,88,91^, which are less tightly anchored to stable anatomical landmarks and are instead defined by distributed gradients of abstraction and task demands. We find that face- and word-selective patches emerge reliably in adults near specific *PdPs*, despite robust VTC selectivity in the same children, underscoring delayed prefrontal maturation. The quadruplet IFS pleating pattern, likely human-specific and under strong genetic control^56^, aligns with spatial layout of category-selective patches, suggesting that frontal sulcal anatomy provides a developmental scaffold for the positioning and refinement of prefrontal functional fields.

### Microstructural Development

Quantitative MRI revealed pruning-like reductions in VLPFC tissue microstructure during adolescence followed by partial recovery in adulthood, a pattern not previously demonstrated non-invasively in living human cortex. This diverges from the steady tissue proliferation observed in VTC^16^ **(Fig. 5)** and argues against a model in which prefrontal maturation simply reflects late-arriving myelinated sensory inputs. Left-right asymmetries in tissue trajectories are consistent with earlier functional reorganization in left VLPFC, potentially reflecting genetic bias supporting language lateralization^99,100^. Developmental gene expression analyses corroborate these qMRI findings, with myelin-associated gene expression recovering post-adolescence while synaptic gene expression remains suppressed, suggesting that pruning disproportionately removed non-myelinated structures. Future work should examine whether this pruning-recovery trajectory is unique to higher-order association cortex or reflects a universal principle of cortical development occurring earlier in sensory regions.

### Broader Implications

These findings advance a conceptual shift in our understanding of prefrontal cortex. Rather than serving solely as a domain-general controller or constructing category geometry de novo, VLPFC organizes early-available category signals into hemisphere-specific taxonomies that are gradually stabilized and geometrically balanced across development. Posterior-anterior coupling strengthens not because prefrontal cortex acquires sensory cortex structure, but because intrinsic prefrontal geometry is reshaped to support selective long-range coordination. Juxtaposed with the early stabilization of VTC, the extended reorganization of VLPFC highlights a core developmental distinction between sensory and prefrontal representational systems, and points toward a broader principle: that human PFC is uniquely positioned to integrate early sensory-driven inputs with anatomically and developmentally constrained scaffolds, producing structured representations that underpin flexible cognition.

## Methods

### Participants

Participants who underwent structural and functional MRI (Figures 1-3) included thirty children (ages 5–12, mean 9.17 ± 2.20, 18 females) and thirty young adults (ages 20–29, mean 23.80 ± 1.79, 22 females). All participants had normal or corrected-to-normal vision. Participants, or their parents, gave written informed consent, children assented to research, and all procedures were approved by the Princeton Internal Review Board on Human Subjects Research.

Tissue dataset: Additional analysis of cortical tissue was conducted in an independent set of participants from a large quantitative MRI dataset discussed in Yeatman et al^69^. These participants included individuals from 5 to 82 years of age (N = 109). For the purposes of our study, older adults ages 55 to 82 (N = 25) were excluded, given that the process of aging is distinct from development. Of the 84 subjects under 55, two were excluded due to missing structural data. The final 82 subjects were separated into four groups: kids ages 5 to 10 years old (N = 30, M = 8.73 ± .87, 12 females), teens ages 11 to 17 years old (N = 18, M = 13.67 ± 2.03, 10 females), young adults ages 18 to 24 years old (N = 11, M = 21.27 ± 1.74, 7 females), and adults ages 25 to 54 years old (N = 23, M = 36.70 ± 8.03, 10 females).

### Data Acquisition

Quantitative magnetic resonance imaging (qMRI): QMRI measurements were obtained using protocols from Mezer et al.^94^. For the larger qMRI dataset presented in Figure 4, four spoiled gradient echo (spoiled-GE) images were acquired to measure T1 relaxation times with flip angles of 4, 10, 20, and 30 degrees (TR = 14ms,TE = 2.4 ms, 0.8 mm x 0.8mm x 1.0 mm). To correct for field inhomogeneities, four additional spin echo inversion recovery scans (SEIR) were performed. These scans utilized an echo planar imaging (EPI) read-out, slab inversion pulse, and spectral spatial fat suppression (TR = 3.0 s, TE = minimum full value). A 2x acceleration was applied during acquisition. Inversion times of 50, 4000, 1200, and 2400 ms were used. The SEIR scans had an in-plane resolution 2.0 mm x 2.0m and a slice thickness of 4.0 mm.Using the quantitative measurements obtained from the spoiled-GE and SEIR scans, an artificial T1-weighted anatomy optimized for tissue segmentation was generated for each participant and used to reconstruct individual cortical surfaces. A second qMRI dataset was also collected using identical parameters on a separate group of participants from the larger dataset^68^. These participants were the same children and adults that completed the functional localizer discussed below. All qMRI data was analyzed with the identical mrQ analysis pipeline: https://github.com/mezera/mrQ.

Functional MRI: Participants underwent 3-Tesla magnetic resonance imaging (General Electric Discovery MR750 3T scanner at Stanford University, or a Siemens 3T MAGNETOM Prisma scanner located in the Scully Center at Princeton University). A phase-array 32-channel head coil was used for the category localizer experiment. Functional data were collected with a simultaneous multislice echo planar imaging (EPI) sequence and a multiplexing factor of 3 to acquire near whole-brain volumes (48 slices, TR = 1s, TE = 30ms). Slices were aligned parallel to the parieto-occipital sulcus at a resolution of 2.4mm isotropic voxels with a T2*-sensitive gradient echo sequence.

Category localizer: Participants underwent an fMRI category localizer experiment, which identifies voxels whose neural response prefers a particular category^101^. Each participant completed 3 runs, each 5 minutes and 18 seconds long. In each run, participants were presented with stimuli from five categories, each with two subcategories (faces: child, adult; bodies: whole, limbs; places: corridors, houses; objects: cars, guitars; characters: words, numbers). Images within the same category were presented in 4 s blocks at a rate of 2 Hz and were not repeated across blocks or runs. 4 s blank trials were also presented throughout a block. During a run, each category was presented eight times in counterbalanced order, with the order differing for each run. Throughout the experiment, participants fixated on a central dot and performed an oddball detection task, pressing a button when phase-scrambled images randomly appeared.

### Data Analysis

Anatomical data analysis: The SEIR and spoiled-GE scans were processed using the mrQ software package in MATLAB to produce the T1-weighted maps. The mrQ analysis pipeline corrected for RF coil bias using SEIR-EPI scans, resulting in accurate macromolecular tissue volume (MTV) and R1 (1/T1) fits across the brain. The complete analysis pipeline and its description can be found at https://github.com/mezera/mrQ. Brain-wide maps of MTV represent the volume of non-water tissue within a given voxel (1 - water fraction), while maps of R1 reflect the average rate at which protons relax to field aligned (1 / T1-relaxation time), and is more sensitive to the composition of tissue within a voxel. Voxels within regions of interest defined along the cortical ribbon were averaged within each participant’s native brain to produce a single tissue metric (average R1 or average MTV) per ROI, per subject. Regions of interest were drawn on the cortical surface, projected across the cortical depth to fill the cortical ribbon, and then brought into the qMRI data space for quantification. The VLPFC and VTC ROI’s encompass all of the sub-regions illustrated in Figure 1F-G. To reduce inter-subject noise, participants are binned within developmental age-groups: children 5-10 years old, teens 11-17, young adults 18-24, adults 25-55. From the quantitative MRI data, a synthetic T1-weighted anatomical image was produced for each individual and used for cortical surface reconstruction and functional data visualization. Artificial T1-weighted images were automatically segmented using FreeSurfer^102^ (FreeSurfer 7.0.0, http://surfer.nmr.mgh.harvard.edu) and then manually validated and corrected using ITKGray to rectify any classification errors. Cortical reconstructions were generated from these segmentations using FreeSurfer 7.0.0. For **Figure 1 D-E**, we quantified the distance of each sulcal pleat (the pair of an annectant gyrus and its neighboring annectant sulcus) to an anatomical reference point that could be readily identified within each individual. Within each participant, we identified the inferior frontal junction as the vertex where the center of the inferior frontal sulcus’ fundus collides with the precentral sulcus. From this vertex, we measured the cortical distance to the center of the four annectant gyri within the IFS using FreeSurfer’s mris_pmake function.

fMRI data analysis: Functional data were motion-corrected within and between scans using the Human Connectome Project (HCP) pipeline and FSL 6.0.4. Data were then processed and analyzed using the Freesurfer Functional Analysis Stream (FSFAST) pipeline. All data were analyzed within the individual participant native brain space. Functional data were automatically aligned to the artificial T1-weighted volume produced from qMRI without smoothing, and were restricted to the cortical ribbon. Contrast maps of faces, pseudowords, bodies, objects, and places versus all other categories were produced for each participant, and where relevant, were thresholded at t-values > 2.5 in the contrast of preferred > non-preferred stimuli. That is, sub-categories (e.g., child and adult faces) were combined within a single contrast. Maps shown in **Fig 1A** were produced by using surface-based alignment to register each category contrast map from native participant space into the shared fsaverage brain space to produce average selectivity maps for each category, in each age group. Functional data represented in **Figures 1B-C, 2, 3A-B, 3E-F, and 4A-B** were all extracted from each participant’s native anatomical space. T-value for a given contrast map were averaged within a given ROI, hand-drawn according to anatomy in each participant’s native space. For **Figure 2**, the mean response within each ROI to a given category was concatenated with the mean responses to other categories individually for each group. These concatenated vectors were then compared using a Pearson correlation whose results are reported in the corner of each panel within **Figure 2**. Functional data represented in **Fig 3C-D** and **Fig 3G-H** were extracted from fsaverage-space contrast maps at vertices defined by each ROI label as the mean across age groups for each hemisphere and contrast. Functional data represented in **Fig 3A–B** were extracted from fsaverage-space contrast maps at vertices defined by each ROI label, separately for each hemisphere, subject, contrast, and group, and organized into ROI-specific feature matrices. The mean voxel pattern for each category within an ROI was calculated by averaging across all voxels for each participant and then across participants within that category. For each ROI, mean voxel patterns per category were clustered using hierarchical clustering to define groups of similar categories, and these cluster labels were then used as targets in a leave-one-subject-out linear SVC to decode category clusters from participant voxel patterns. Decoding performance was quantified as classification accuracy, and shuffled empirical chance was computed by repeating the same decoding procedure on 1000 randomly permuted labels to estimate chance-level performance.

Gene expression analysis: The data reported in **Figure 4F** were obtained from the Allen Human Brain Atlas, in particular the BrainSpan Atlas of the Developing Human Brain and its Developmental Transcriptome. These resources are freely available at the following hyperlink: https://www.brainspan.org/. Exon microarray and associated metadata for the RNA-seq analyses summarized to genes were acquired, which samples over 17,000 genes from gestational to 40 years of age. We included data from the ventrolateral frontal cortex (VFC) region within the dataset because this tissue location was centered on the posterior inferior frontal gyrus, which is encompassed within the VLPFC ROI used within our functional and quantitative MRI data. Therefore, there is strong spatial correspondence between the gene expression and MRI data, which affords stronger comparison of these different datatypes. To identify the genes related to synaptic structures, we acquired a synaptic gene ontology set list through the SynGo database^61^ (https://syngoportal.org/) which identifies, through consensus of an international research community, those genes localized to, and involved in, synaptic structures of neural tissue. For myelin-related structures, we used genes that strongly co-localize and are known to be directly involved in the structure and maintenance of human myelin sheath^62^: PLP1, PLLP, MAG, MBP, CA2, PMP22, MAL, MBP, ERMN, OMG. Microarray expression data acquired from the human brain atlas have been preprocessed and normalized per their published pipelines. After intersecting the full developmental transcriptome gene set with our myelin and synapse gene sets of interest, we further normalized the expression of each gene to the average expression level during adolescence (11<age in years<20) within a set, before averaging across all genes within each set. Average expression levels from ages 11 to 40 years of age are then plotted in **Figure 4F**.

### Definitions of regions of interest

VLPFC: Ventrolateral prefrontal cortex (VLPFC) is defined as in Levy and Wagner^23^ and Bugatus et al.^98^. VLPFC is posteriorly bounded by the caudal lip of the precentral sulcus and the ascending ramus of the lateral fissure (aalf). The anterior boundary lies on the most anterior edge of the pars orbitalis. The superior boundary lies on the dorsal lip of the IFS, and the inferior boundary lies on the dorsal lip of the lateral fissure. Because the frontal eye field (FEF) is heavily implicated in eye movements and viewing visual stimuli^101^, we also include the inferior portion of the superior precentral sulcus, below the superior frontal sulcus, in our region of interest. An example of the VLPFC ROI can be seen in **Figure 1**. To achieve finer-grained analysis of the frontal lobe, we parcellated VLPFC into 10 ROIs in every individual **(Fig 1D)**. The IFS was divided into four ROIs along its anterior-posterior axis according to *plis de passage* (e.g., cortical pleats) observed consistently across participants. Each IFS subsection included one pleat: (1) the inferior frontal junction (IFJ), which extends anteriorly from the PCS to include PdP1 into the sulcal bed just anterior to it, (2) the posterior IFS (pIFS), which extends from the IFJ ROI to the sulcal bed of PdP2, (3) the middle IFS (mIFS), which begins at the anterior border of the pIFS and ends at the sulcal bed of PdP3, and (4) the anterior IFS (aIFS), which starts at the mIFS and extends to the end of the IFS near the lateral frontomarginal sulcus (lfms). The inferior precentral sulcus was divided into superior and inferior components (s–IPCS, i–IPCS) separated by another *PdP* which extends towards the IFJ. The inferior portion of the superior precentral sulcus and the surrounding gyrus were included and referred to here as the SPCS ROI. We also included the op, tr, and or – which subdivide the inferior frontal gyrus – as three separate ROIs^61,102^.

VTC: To define VTC, we combine cytoarchitectonic regions FG1, FG2, FG3, FG4, hOc3v, hOc4v, hO3cvla, and hO4cvlp **(Fig 1E)**. These cytoarchitectonic regions were defined original postmortem brains, which were then resampled to cortical surface reconstructions using FreeSurfer^62^. Anatomically, we follow the boundaries proposed in Rosenke et al. 2018^62^ which includes a lateral boundary at the occipitotemporal sulcus, a medial boundary at the parahippocampal gyrus, and an anterior boundary at the tip of the mid-fusiform sulcus. However, we extend VTC beyond the posterior transverse collateral sulcus, which is the posterior anatomical boundary, so that it includes cytoarchitectonic regions hOc3v, hOc4v, hO3cvla, and hO4cvlp. In our subsequent finer-grained analysis comparing VTC and VLPFC, this allows for comparison using a similar number of smaller ROIs within each lobe. Furthermore, cytoarchitectonic regions largely follow cortical folding patterns, making them further comparable to the ROI’s drawn within VLPFC which were drawn relative to cortical folding patterns.

## Data and Code Availability

All data and analysis code used in this study are publicly available at GitHub: https://github.com/priscillalouis. The repository includes scripts required to reproduce the main figures and results.

## Statistical Information

All statistical analyses were conducted in MATLAB and Python. Group-level effects were assessed using t-tests and ANOVA models. For between-group comparisons (adults vs children), independent-samples t-tests were used, and within-subject effects were assessed using paired-samples t-tests. One-sample t-tests were used to evaluate whether activation or distance measures differed from zero or chance where applicable. Multi-factor effects were assessed using factorial ANOVAs and repeated-measures ANOVAs. Where applicable, F-statistics, degrees of freedom, and p-values are reported for ANOVAs, and t-statistics with corresponding degrees of freedom are reported for t-tests. Effect sizes were quantified using partial eta-squared for ANOVA models and Cohen’s d for t-tests (pooled SD for independent samples; *dz* for paired comparisons). Correlation analyses were performed using Pearson correlation coefficients. Multiple comparisons were controlled using false discovery rate (FDR; Benjamini–Hochberg) correction. Error bars in figures reflect either standard error of the mean or subject-wise variability as specified in figure legends, and all analyses report exact sample sizes used for each test, with subject-level values treated as independent replicates.

## Author Contributions

Authors **PL**, **JY**, **ZY**, **RM**, **TB** and **JG** contributed to data analysis, **JY** and **JG** contributed to experimental design, **PL** and **JG** contributed to data collection, and **PL**, **JY** and **JG** contributed to manuscript preparation.

## Acknowledgements

The authors acknowledge the support of the Scully Center at Princeton University and Stanford University for providing access to MRI facilities and for technical support through data acquisition. We are deeply grateful to the families and participants whose time and commitment made this study possible.

## Supplemental Information

**Supplementary Figure 1.**
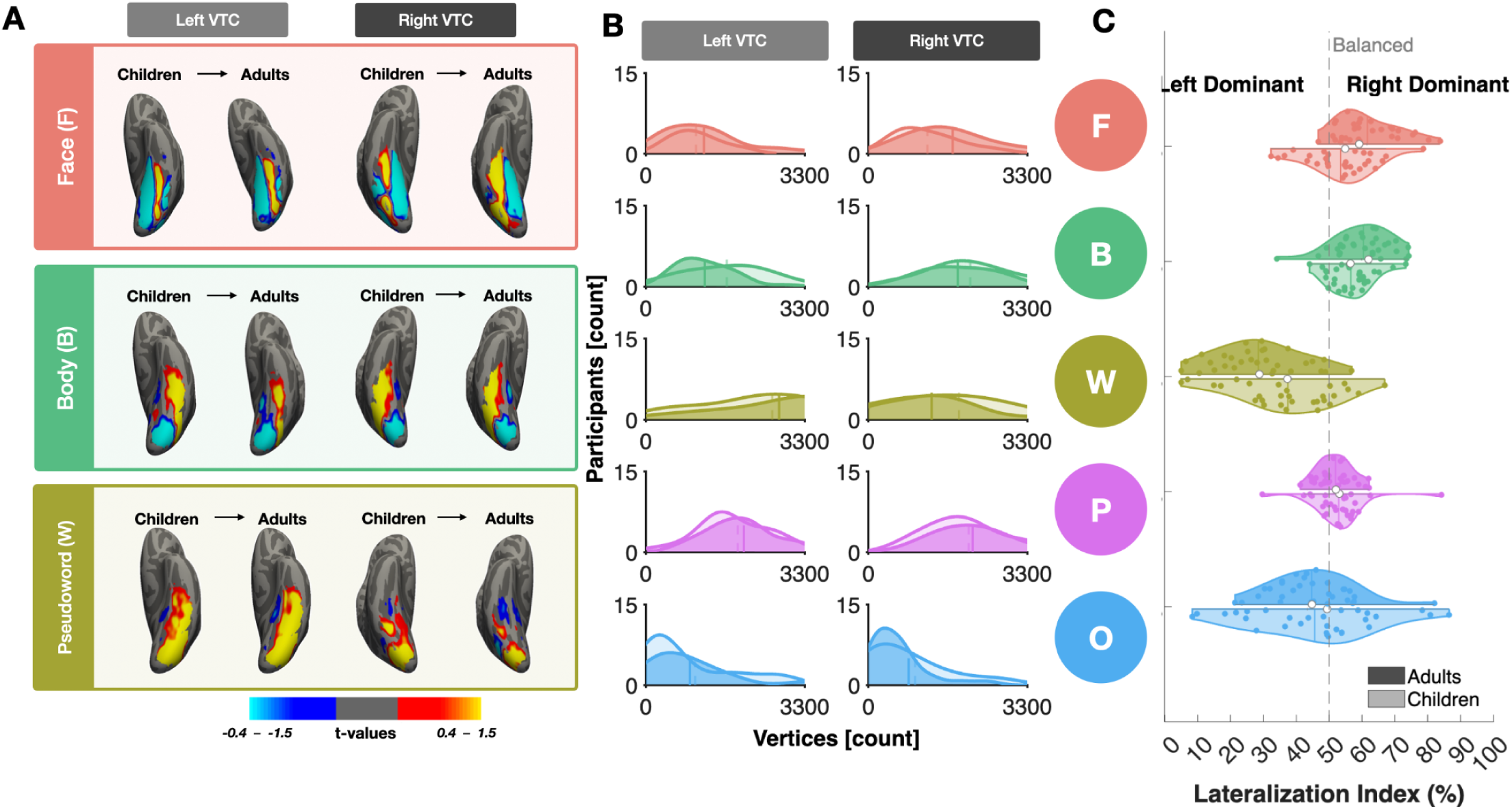
VTC shows robust, canonical category-selective organization. **(A)** Average cortical surface maps of category selectivity (preferred > all others) for faces, bodies, and words on the fsaverage template. Columns show a sample stimulus, left hemisphere maps, and right hemisphere maps; each hemisphere panel shows children (left) and adults (right). Diverging colors indicate positive (red–yellow) and negative (blue) selectivity. **(B)** Density plots of suprathreshold vertex counts (t-values > 2.5) for five visual categories in each hemisphere; translucent and solid curves indicate children and adults, respectively; vertical lines show group means. **(C)** Split-violin plots of hemispheric dominance (% RH activation relative to LH + RH) per category for children (lighter) and adults (darker); values above and below 50% indicate RH and LH dominance, respectively.

**Supplementary Figure 2.**
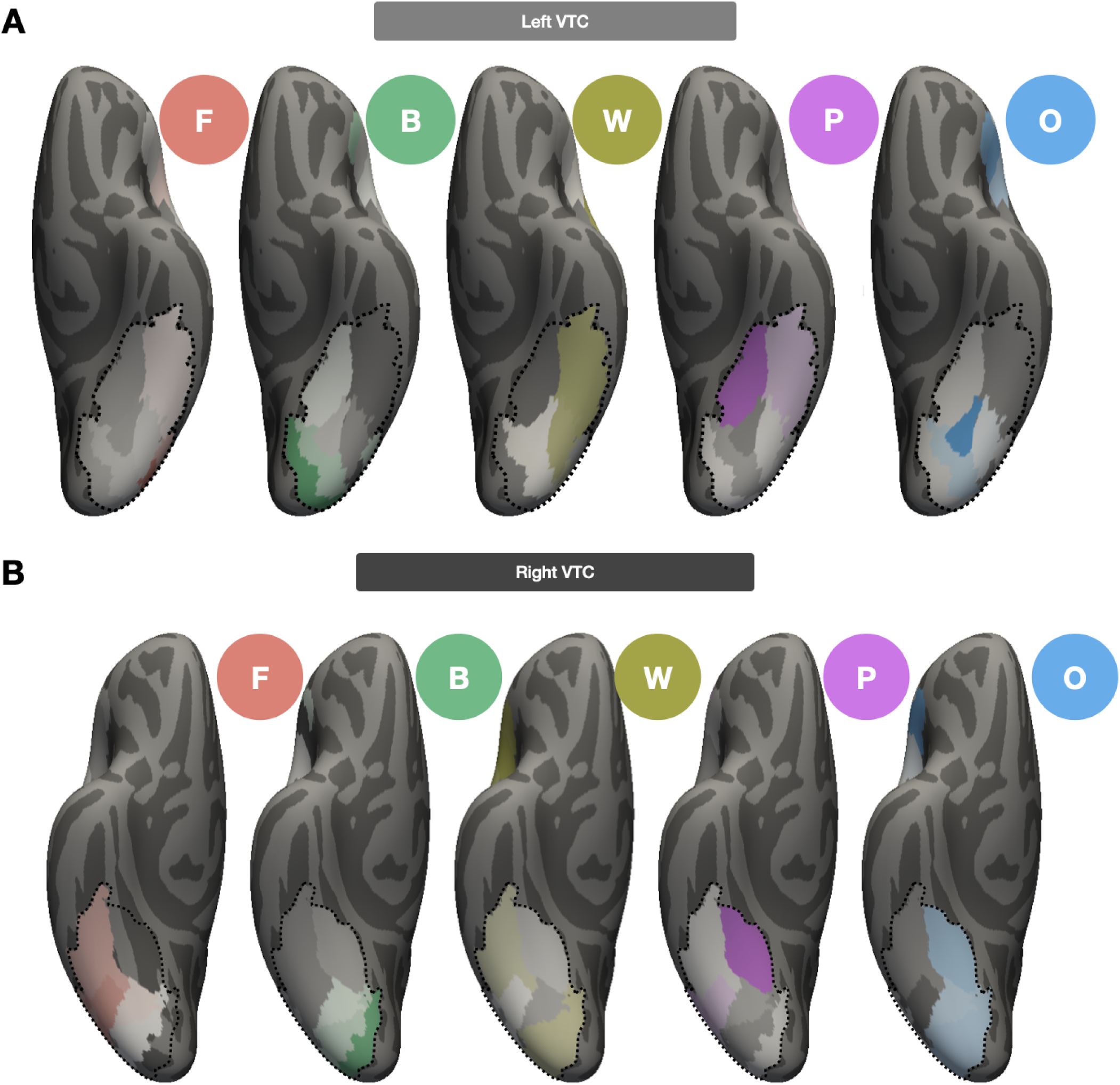
VTC category response patterns show modest developmental differences relative to VLPFC. **(A)** Z-scored developmental difference maps (adults minus children, normalized within each system) projected onto LH VTC surface. Positive/negative values indicate stronger adult/child responses. **(B)** Same as A for RH VTC.

**Supplementary Figure 3.**
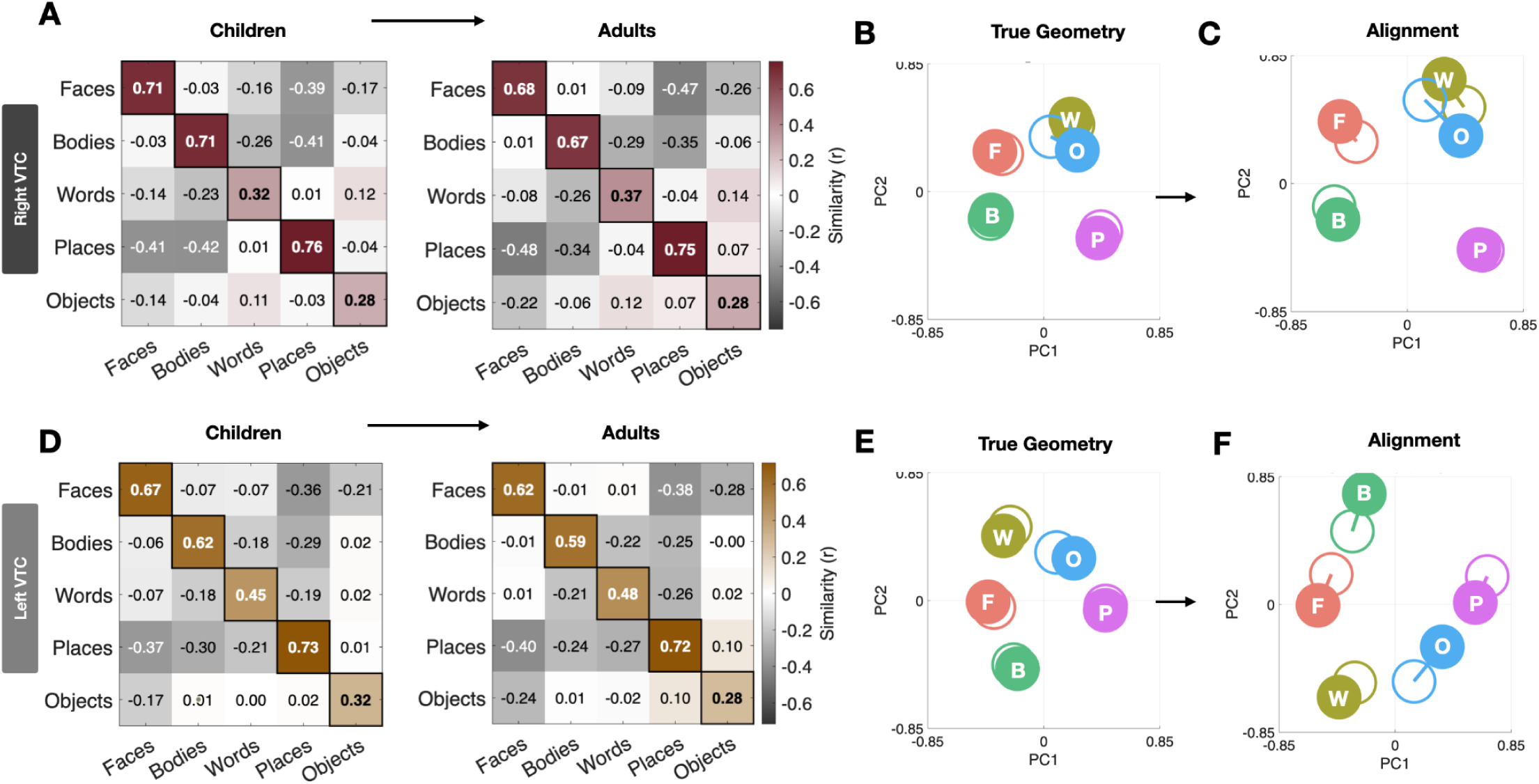
VTC representational geometry is stable across development. **(A)** LH VTC representational similarity matrices (mean pairwise Pearson r between voxelwise t-value maps for five categories) computed using LORO cross-validation, shown separately for children and adults. Diagonal = within-category reliability; off-diagonal = cross-category similarity. **(B)** PCA of voxelwise category-selective response patterns in LH VTC, plotted in PC1–PC2 space (z-scored across vertices). Filled/open circles = adults/children; lines connect matched categories across groups. **(C)** Same as B, following Procrustes alignment of children to the adult configuration. **(D–F)** Same as A–C for RH VTC.

## References

1. Kanwisher, N., McDermott, J. & Chun, M. M. The Fusiform Face Area: A Module in Human Extrastriate Cortex Specialized for Face Perception. J. Neurosci. 17, 4302–4311 (1997).

2. Golarai, G. et al. Differential development of high-level visual cortex correlates with category-specific recognition memory. Nat. Neurosci. 10, 512–522 (2007).

3. Scherf, K. S., Behrmann, M., Humphreys, K. & Luna, B. Visual category-selectivity for faces, places and objects emerges along different developmental trajectories. Dev. Sci. 10, F15–30 (2007).

4. Liu, C. et al. The Visual Word Form Area: evidence from an fMRI study of implicit processing of Chinese characters. NeuroImage 40, 1350–1361 (2008).

5. McCandliss, B. D., Cohen, L. & Dehaene, S. The visual word form area: expertise for reading in the fusiform gyrus. Trends Cogn. Sci. 7, 293–299 (2003).

6. Dehaene-Lambertz, G., Monzalvo, K. & Dehaene, S. The emergence of the visual word form: Longitudinal evolution of category-specific ventral visual areas during reading acquisition. PLOS Biol. 16, e2004103 (2018).

7. Dehaene, S. et al. How learning to read changes the cortical networks for vision and language. Science 330, 1359–1364 (2010).

8. Arcaro, M. J., Schade, P. F., Vincent, J. L., Ponce, C. R. & Livingstone, M. S. Seeing faces is necessary for face-domain formation. Nat. Neurosci. 20, 1404–1412 (2017).

9. Gomez, J., Barnett, M. & Grill-Spector, K. Extensive childhood experience with Pokémon suggests eccentricity drives organization of visual cortex. *Nat*. Hum. Behav. 3, 611–624 (2019).

10. Weiner, K. S. & Grill-Spector, K. Neural representations of faces and limbs neighbor in human high-level visual cortex: evidence for a new organization principle. Psychol. Res. 77, 74–97 (2013).

11. Weiner, K. S. et al. The mid-fusiform sulcus: a landmark identifying both cytoarchitectonic and functional divisions of human ventral temporal cortex. NeuroImage 84, 453–465 (2014).

12. Weiner, K. S. et al. Defining the most probable location of the parahippocampal place area using cortex-based alignment and cross-validation. NeuroImage 170, 373–384 (2018).

13. Nordt, M. et al. Cortical recycling in high-level visual cortex during childhood development. *Nat*. Hum. Behav. 5, 1686–1697 (2021).

14. Op de Beeck, H. P., Pillet, I. & Ritchie, J. B. Factors Determining Where Category-Selective Areas Emerge in Visual Cortex. Trends Cogn. Sci. 23, 784–797 (2019).

15. Grill-Spector, K. & Weiner, K. S. The functional architecture of the ventral temporal cortex and its role in categorization. Nat. Rev. Neurosci. 15, 536–548 (2014).

16. Gomez, J. et al. Microstructural proliferation in human cortex is coupled with the development of face processing. Science 355, 68–71 (2017).

17. Srihasam, K., Vincent, J. L. & Livingstone, M. S. Novel domain formation reveals proto-architecture in inferotemporal cortex. Nat. Neurosci. 17, 1776–1783 (2014).

18. Nordt, M. et al. Learning to Read Increases the Informativeness of Distributed Ventral Temporal Responses. Cereb. Cortex 29, 3124–3139 (2019).

19. Golarai, G., Liberman, A. & Grill-Spector, K. Experience Shapes the Development of Neural Substrates of Face Processing in Human Ventral Temporal Cortex. Cereb. Cortex N. Y. N 1991 27, 1229–1244 (2017).

20. Chafee, M. V. & Heilbronner, S. R. Prefrontal cortex. Curr. Biol. CB 32, R346–R351 (2022).

21. Bugatus, L., Weiner, K. S. & Grill-Spector, K. Task alters category representations in prefrontal but not high-level visual cortex. NeuroImage 155, 437–449 (2017).

22. Friedman, N. P. & Robbins, T. W. The role of prefrontal cortex in cognitive control and executive function. Neuropsychopharmacology 47, 72–89 (2022).

23. Levy, B. J. & Wagner, A. D. Cognitive control and right ventrolateral prefrontal cortex: reflexive reorienting, motor inhibition, and action updating. Ann. N. Y. Acad. Sci. 1224, 40–62 (2011).

24. Costantini, I. et al. A cellular resolution atlas of Broca’s area. Sci. Adv. 9, eadg3844 (2023).

25. Amunts, K. & Zilles, K. Architecture and organizational principles of Broca’s region. Trends Cogn. Sci. 16, 418–426 (2012).

26. Sprung-Much, T., Eichert, N., Nolan, E. & Petrides, M. Broca’s area and the search for anatomical asymmetry: commentary and perspectives. Brain Struct. Funct. 227, 441–449 (2022).

27. Chan, A. W.-Y. & Downing, P. E. Faces and eyes in human lateral prefrontal cortex. Front. Hum. Neurosci. 5, 51 (2011).

28. Tsao, D. Y., Moeller, S. & Freiwald, W. A. Comparing face patch systems in macaques and humans. Proc. Natl. Acad. Sci. 105, 19514–19519 (2008).

29. Nikel, L., Sliwinska, M. W., Kucuk, E., Ungerleider, L. G. & Pitcher, D. Measuring the response to visually presented faces in the human lateral prefrontal cortex. Cereb. Cortex Commun. 3, tgac036 (2022).

30. Stone, H. L. et al. Anatomically Distinct Regions in the Inferior Frontal Cortex Are Modulated by Task and Reading Skill. J. Neurosci. 45, (2025).

31. Wagner, A. D., Gabrieli, J. D. & Verfaellie, M. Dissociations between familiarity processes in explicit recognition and implicit perceptual memory. J. Exp. Psychol. Learn. Mem. Cogn. 23, 305–323 (1997).

32. Clark, D. & Wagner, A. D. Assembling and encoding word representations: fMRI subsequent memory effects implicate a role for phonological control. Neuropsychologia 41, 304–317 (2003).

33. Dumoulin, S. O. et al. A new anatomical landmark for reliable identification of human area V5/MT: a quantitative analysis of sulcal patterning. Cereb. Cortex N. Y. N 1991 10, 454–463 (2000).

34. Weiner, K. S. & Grill-Spector, K. Not one extrastriate body area: Using anatomical landmarks, hMT+, and visual field maps to parcellate limb-selective activations in human lateral occipitotemporal cortex. NeuroImage 56, 2183–2199 (2011).

35. Witthoft, N. et al. Where is human V4? Predicting the location of hV4 and VO1 from cortical folding. Cereb. Cortex N. Y. N 1991 24, 2401–2408 (2014).

36. Benson, N. C. & Winawer, J. Bayesian analysis of retinotopic maps. eLife 7, e40224 (2018).

37. Gomez, J. et al. Development of population receptive fields in the lateral visual stream improves spatial coding amid stable structural-functional coupling. NeuroImage 188, 59–69 (2019).

38. Sanides, F. Structure and function of the human frontal lobe. Neuropsychologia 2, 209–219 (1964).

39. Voorhies, W. I., Miller, J. A., Yao, J. K., Bunge, S. A. & Weiner, K. S. Cognitive insights from tertiary sulci in prefrontal cortex. Nat. Commun. 12, 5122 (2021).

40. Willbrand, E. H., Voorhies, W. I., Yao, J. K., Weiner, K. S. & Bunge, S. A. Presence or absence of a prefrontal sulcus is linked to reasoning performance during child development. Brain Struct. Funct. 227, 2543–2551 (2022).

41. Willbrand, E. H., Ferrer, E., Bunge, S. A. & Weiner, K. S. Development of Human Lateral Prefrontal Sulcal Morphology and Its Relation to Reasoning Performance. J. Neurosci. 43, 2552–2567 (2023).

42. Tissier, C. et al. Sulcal Polymorphisms of the IFC and ACC Contribute to Inhibitory Control Variability in Children and Adults. eNeuro 5, (2018).

43. Parker, P. R. L. et al. A dynamic sequence of visual processing initiated by gaze shifts. Nat. Neurosci. 26, 2192–2202 (2023).

44. Petit, L. & Pouget, P. The comparative anatomy of frontal eye fields in primates. Cortex J. Devoted Study Nerv. Syst. Behav. 118, 51–64 (2019).

45. Rakic, P., Bourgeois, J. P., Eckenhoff, M. F., Zecevic, N. & Goldman-Rakic, P. S. Concurrent overproduction of synapses in diverse regions of the primate cerebral cortex. Science 232, 232–235 (1986).

46. Huttenlocher, P. R. & Dabholkar, A. S. Regional differences in synaptogenesis in human cerebral cortex. J. Comp. Neurol. 387, 167–178 (1997).

47. Elston, G. N. & Fujita, I. Pyramidal cell development: postnatal spinogenesis, dendritic growth, axon growth, and electrophysiology. Front. Neuroanat. 8, (2014).

48. Bugatus, L., Weiner, K. & Grill-Spector, K. Task modulates category selectivity along a gradient from occipitotemporal cortex to prefrontal cortex in word- and face-selective regions. J. Vis. 15, 1170–1170 (2015).

49. Maurer, U., Rossion, B. & McCandliss, B. D. Category Specificity in Early Perception: Face and Word N170 Responses Differ in Both Lateralization and Habituation Properties. Front. Hum. Neurosci. 2, 18 (2008).

50. Rossion, B. Constraining the cortical face network by neuroimaging studies of acquired prosopagnosia. NeuroImage 40, 423–426 (2008).

51. Dundas, E. M., Plaut, D. C. & Behrmann, M. The joint development of hemispheric lateralization for words and faces. J. Exp. Psychol. Gen. 142, 348–358 (2013).

52. Dundas, E. M., Plaut, D. C. & Behrmann, M. An ERP investigation of the co-development of hemispheric lateralization of face and word recognition. Neuropsychologia 61, 315–323 (2014).

53. Sacchi, E. & Laszlo, S. An event-related potential study of the relationship between N170 lateralization and phonological awareness in developing readers. Neuropsychologia 91, 415–425 (2016).

54. Malik-Moraleda, S. et al. An investigation across 45 languages and 12 language families reveals a universal language network. Nat. Neurosci. 25, 1014–1019 (2022).

55. Gratiolet, P. L. & Royal College of Surgeons of England. Mémoire sur les plis cérébraux de l’homme et des primatès. (Paris : Arthus Bertrand, 1854).

56. Mangin, J.-F. et al. “Plis de passage” Deserve a Role in Models of the Cortical Folding Process. Brain Topogr. 32, 1035–1048 (2019).

57. Simone, L. et al. Distinct Functional and Structural Connectivity of the Human Hand-Knob Supported by Intraoperative Findings. J. Neurosci. 41, 4223–4233 (2021).

58. Muellen, A. M. & Schweizer, R. Characterization of the Central Sulcus Pli-De-Passage Fronto-Pariétal Moyen in > 1000 Human Brains. Hum. Brain Mapp. 47, e70457 (2026).

59. Im, K. & Grant, P. E. Sulcal pits and patterns in developing human brains. NeuroImage 185, 881–890 (2019).

60. Bodin, C. et al. Plis de passage in the superior temporal sulcus: Morphology and local connectivity. NeuroImage 225, 117513 (2021).

61. Atlas of the Morphology of the Human Cerebral Cortex on the Average MNI Brain - Edition 1 - By Michael Petrides Elsevier Health Inspection Copies. https://www.inspectioncopy.elsevier.com/book/details/9780128009321.

62. Rosenke, M. et al. A cross-validated cytoarchitectonic atlas of the human ventral visual stream. NeuroImage 170, 257–270 (2018).

63. Ritchie, J. B. et al. Untangling the Animacy Organization of Occipitotemporal Cortex. J. Neurosci. 41, 7103–7119 (2021).

64. Wiggett, A. J., Pritchard, I. C. & Downing, P. E. Animate and inanimate objects in human visual cortex: Evidence for task-independent category effects. Neuropsychologia 47, 3111–3117 (2009).

65. Abdel-Ghaffar, S. A. et al. Occipital-temporal cortical tuning to semantic and affective features of natural images predicts associated behavioral responses. Nat. Commun. 15, 5531 (2024).

66. Abdallah, M., Zanitti, G. E., Iovene, V. & Wassermann, D. Functional gradients in the human lateral prefrontal cortex revealed by a comprehensive coordinate-based meta-analysis. eLife 11, e76926 (2022).

67. van Blooijs, D. et al. Developmental trajectory of transmission speed in the human brain. Nat. Neurosci. 26, 537–541 (2023).

68. Liao, Z. et al. Hemispheric asymmetry in cortical thinning reflects intrinsic organization of the neurotransmitter systems and homotopic functional connectivity. Proc. Natl. Acad. Sci. U. S. A. 120, e2306990120 (2023).

69. Yeatman, J. D., Wandell, B. A. & Mezer, A. A. Lifespan maturation and degeneration of human brain white matter. Nat. Commun. 5, 4932 (2014).

70. Giallanza, T., Campbell, D., Cohen, J. D. & Rogers, T. T. An integrated model of semantics and control. Psychol. Rev. 132, 1128–1177 (2025).

71. Miller, E. K. & Cohen, J. D. An integrative theory of prefrontal cortex function. Annu. Rev. Neurosci. 24, 167–202 (2001).

72. Broschard, M. B., Turner, B. M., Tranel, D. & Freeman, J. H. Dissociable Roles of the Dorsolateral and Ventromedial Prefrontal Cortex in Human Categorization. J. Neurosci. Off. J. Soc. Neurosci. 44, e2343232024 (2024).

73. Stigliani, A., Weiner, K. S. & Grill-Spector, K. Temporal Processing Capacity in High-Level Visual Cortex Is Domain Specific. J. Neurosci. 35, 12412–12424 (2015).

74. Chan, A. W. Y. Functional organization and visual representations of human ventral lateral prefrontal cortex. Front. Psychol. 4, (2013).

75. Duan, Y., Zhan, J., Gross, J., Ince, R. A. A. & Schyns, P. G. Pre-frontal cortex guides dimension-reducing transformations in the occipito-ventral pathway for categorization behaviors. Curr. Biol. 34, 3392–3404.e5 (2024).

76. Erez, Y. & Duncan, J. Discrimination of Visual Categories Based on Behavioral Relevance in Widespread Regions of Frontoparietal Cortex. J. Neurosci. 35, 12383–12393 (2015).

77. Freedman, D. J., Riesenhuber, M., Poggio, T. & Miller, E. K. Categorical Representation of Visual Stimuli in the Primate Prefrontal Cortex. Science 291, 312–316 (2001).

78. Mackey, W. E., Winawer, J. & Curtis, C. E. Visual field map clusters in human frontoparietal cortex. eLife 6, e22974 (2017).

79. Merchant, H., Crowe, D. A., Robertson, M., Fortes, A. F. & Georgopoulos, A. P. Top-Down Spatial Categorization Signal from Prefrontal to Posterior Parietal Cortex in the Primate. Front. Syst. Neurosci. 5, (2011).

80. Rajimehr, R., Young, J. C. & Tootell, R. B. H. An anterior temporal face patch in human cortex, predicted by macaque maps. Proc. Natl. Acad. Sci. 106, 1995–2000 (2009).

81. Tsao, D. Y., Schweers, N., Moeller, S. & Freiwald, W. A. Patches of face-selective cortex in the macaque frontal lobe. Nat. Neurosci. 11, 877–879 (2008).

82. Badre, D. What Is the Nature of the Hierarchical Organization of Lateral Prefrontal Cortex? Evidence for Homology Across Species. in The Frontal Cortex: Organization, Networks, and Function (eds Banich, M. T., Haber, S. N. & Robbins, T. W.) (MIT Press, Cambridge (MA), 2024).

83. Gonzalez Alam, T. R. del J., et al. A tale of two gradients: differences between the left and right hemispheres predict semantic cognition. Brain Struct. Funct. 227, 631–654 (2022).

84. Ferrante, O. et al. Adversarial testing of global neuronal workspace and integrated information theories of consciousness. Nature 642, 133–142 (2025).

85. He, Y. et al. Diverse Frontoparietal Connectivity Supports Semantic Prediction and Integration in Sentence Comprehension. J. Neurosci. 45, (2025).

86. Keller, A. S. et al. Hierarchical functional system development supports executive function. Trends Cogn. Sci. 27, 160–174 (2023).

87. Reynolds, J. E., Long, X., Grohs, M. N., Dewey, D. & Lebel, C. Structural and functional asymmetry of the language network emerge in early childhood. Dev. Cogn. Neurosci. 39, 100682 (2019).

88. Nee, D. E. & D’Esposito, M. The hierarchical organization of the lateral prefrontal cortex. eLife 5, e12112 (2016).

89. Andrulyte, I. et al. The Relationship between White Matter Architecture and Language Lateralization in the Healthy Brain. J. Neurosci. 44, (2024).

90. Hertrich, I., Dietrich, S., Blum, C. & Ackermann, H. The Role of the Dorsolateral Prefrontal Cortex for Speech and Language Processing. Front. Hum. Neurosci. 15, (2021).

91. Badre, D. & Nee, D. E. Frontal Cortex and the Hierarchical Control of Behavior. Trends Cogn. Sci. 22, 170–188 (2018).

92. Ralph, M. A. L., Jefferies, E., Patterson, K. & Rogers, T. T. The neural and computational bases of semantic cognition. Nat. Rev. Neurosci. 18, 42–55 (2017).

93. Germann, J., Robbins, S., Halsband, U. & Petrides, M. Precentral sulcal complex of the human brain: morphology and statistical probability maps. J. Comp. Neurol. 493, 334–356 (2005).

94. Mezer, A. et al. Quantifying the local tissue volume and composition in individual brains with magnetic resonance imaging. Nat. Med. 19, 1667–1672 (2013).

95. Fischl, B., Sereno, M. I. & Dale, A. M. Cortical Surface-Based Analysis: II: Inflation, Flattening, and a Surface-Based Coordinate System. NeuroImage 9, 195–207 (1999).

96. Koopmans, F. et al. SynGO: An Evidence-Based, Expert-Curated Knowledge Base for the Synapse. Neuron 103, 217–234.e4 (2019).

97. Lee, P. R. & Fields, D. Regulation of myelin genes implicated in psychiatric disorders by functional activity in axons. Front. Neuroanat. 3, (2009).

98. Bugatus, L., Weiner, K. & Grill-Spector, K. Differential representation of category and task information across high level visual cortex and ventro-lateral prefrontal cortex. J. Vis. 16, 256–256 (2016).

99. Witelson, S. F. & Pallie, W. LEFT HEMISPHERE SPECIALIZATION FOR LANGUAGE IN THE NEWBORN: NEUROANATOMICAL EVIDENCE OF ASYMMETRY. Brain 96, 641–646 (1973).

100. Dehaene-Lambertz, G. et al. Functional organization of perisylvian activation during presentation of sentences in preverbal infants. Proc. Natl. Acad. Sci. 103, 14240–14245 (2006).

101. Konen, C. S. & Kastner, S. Representation of Eye Movements and Stimulus Motion in Topographically Organized Areas of Human Posterior Parietal Cortex. J. Neurosci. 28, 8361–8375 (2008).

102. Harvey, D. Y. Neuroanatomy of Language Regions of the Human Brain. J. Undergrad. Neurosci. Educ. 13, R12–R13 (2015).

